# Moderate activity of RNA chaperone maximizes the yield of self-spliced pre-RNA *in vivo*

**DOI:** 10.1101/2022.09.20.508801

**Authors:** Yonghyun Song, D. Thirumalai, Changbong Hyeon

## Abstract

CYT-19 is a DEAD-box protein whose ATP-dependent helicase activity facilitates the folding of group I introns in precursor RNA (pre-RNA) of *Neurospora crassa*. In the process they consume a substantial amount of ATP. While much of the mechanistic insights into CYT-19 activity has been gained through the studies on the folding of *Tetrahymena* group I intron ribozyme, the more biologically relevant issue, namely the effect of CYT-19 on the self-splicing of pre-RNA, remains largely unexplored. Here, we employ a kinetic network model, based on the generalized iterative annealing mechanism, to investigate the relation between CYT-19 activity, rate of ribozyme folding, and the kinetics of the self-splicing reaction. The network rate parameters are extracted by analyzing the recent biochemical data for CYT-19-facilitated folding of *T*. ribozyme. We then build extended models to explore the metabolism of pre-RNA. We show that the timescales of chaperone-mediated folding of group I ribozyme and self-splicing reaction compete with each other. As a consequence, in order to maximize the self-splicing yield of group I introns in pre-RNA, the chaperone activity must be sufficiently large to unfold the misfolded structures, but not too large to unfold the native structures prior to the self-splicing event. We discover that despite the promiscuous action on structured RNAs, the helicase activity of CYT-19 on group I ribozyme gives rise to self-splicing yields that are close to the maximum.

**Significance Statement:** In cells, RNA chaperones assist misfolding-prone ribozymes to fold correctly to carry out its biological function. CYT-19 is an ATP-consuming RNA chaperone that accelerates the production of native group I intron ribozyme by partially unfolding the kinetically trapped structures. Using the theoretical framework based on the iterative annealing mechanism, we establish that to maximize the processing of pre-RNA, an optimal balance should exist between the timescales of self-splicing activity and CYT-19-mediated production of the native ribozyme. Remarkably, the activity of CYT-19 has been optimized to unfold the misfolded structures but is not so high that it disrupts the native ribozyme, which ensures that the yield of the self-splicing reaction is maximized in a biologically relevant time scale.

---

**T**he enzymatic functions that are carried out by structured RNAs (1–5) regulate a diverse array of cellular processes including translation, RNA processing, the maintenance of chromosome ends, and the regulation of gene expression (6–9). However, the homopolymeric nature of the interactions between the nucleotides and the multivalent ion-mediated tertiary interaction result in a highly rugged RNA folding landscapes (4). Even RNA secondary structures are heterogeneous, making interconversions between them extremely slow (10–13). Subsequent transitions of RNA to a functional native structure occur via multiple pathways, usually with diverse timescales, giving rise to multi-exponential kinetics, which is quantitatively explained by the kinetic partitioning mechanism (KPM) (4, 14, 15). Only a small fraction of the ensemble (Φ ≪ 1) folds directly to the native basin of attraction, whereas the remaining fraction (1 − Φ) are trapped in the manifold of long-lived misfolded states that define the competing basins of attraction (Fig. 1) (4). Although spontaneous transitions from misfolded to native states, in principle, are not unattainable, they are kinetically inaccessible because the timescales for such processes are far too long compared to the typical cell doubling time (16–19). RNA chaperones, which assist RNA molecules to escape from deep kinetic traps in the folding landscapes and facilitate their folding on a biologically relevant timescale, are the key players in cellular RNA metabolism (20–23).

**Fig. 1.**
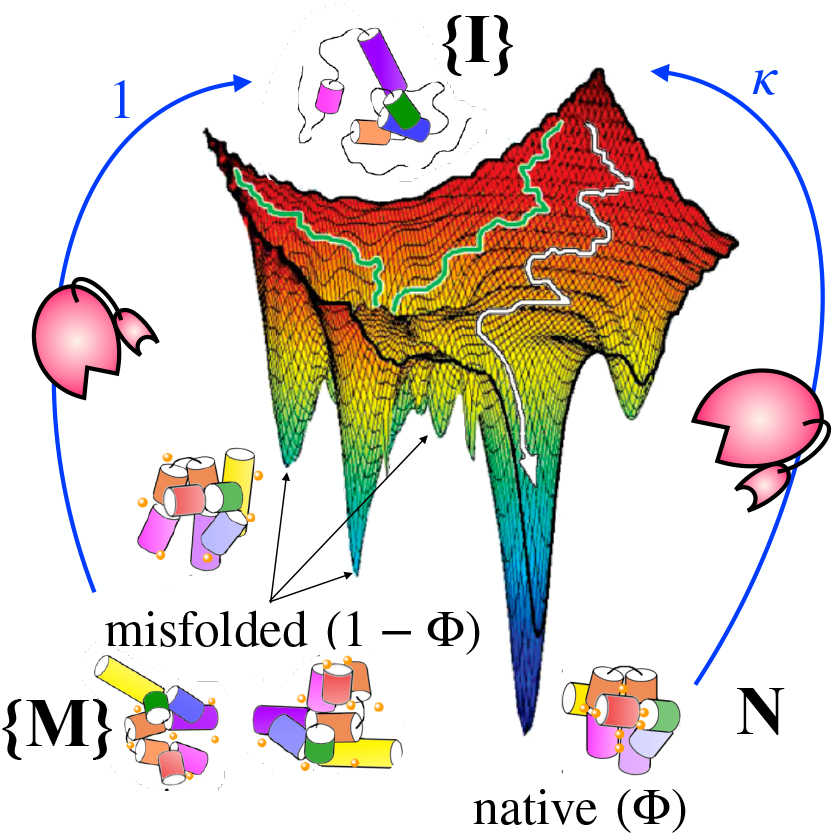
Chaperone-facilitated folding of RNA on a rugged folding landscape. The fraction Φ among the ensemble of RNA molecules in the intermediate (**{I}**) states reaches the native (**N**) state rapidly, while (1 − Φ) are trapped in the ensemble of misfolded (**{M}**) states. The repeated actions of RNA chaperone (depicted as a Pac-Man-like object) anneal the otherwise misfolding-prone population of RNA into the one with a high native yield as quantified in Eq. 1.

CYT-19 in *Neurospora crassa* and its homologue Mss116 in yeast are the best studied RNA chaperones (9, 24). An earlier work showed that unspliced group I introns accumulated in the mutant *N. crassa* that is devoid of CYT-19, a DEAD-box protein with a helicase domain whose ATP-dependent activity accelerates the folding and self-splicing of group I introns (25). Interestingly, although it is a protein derived from fungi, CYT-19 has a generic activity towards RNAs from different organisms (25, 26), as does the bacterial GroEL, the protein chaperone (27, 28).

Russell and colleagues have quantified the effect of CYT-19 on the folding of *T*. ribozyme, a group I intron derived catalytic RNA (29–32), over a wide range of experimental conditions (33–37). They have determined that CYT-19 is generally reactive towards surface-exposed RNA helices (33). Surprisingly, unlike protein chaperones (38), CYT-19 disrupts the structures of both the misfolded and native ribozymes (35, 36).

We have previously provided a mechanistic explanation of the action of RNA chaperones, which consume ATP for their functions, by generalizing (39) the iterative annealing mechanism (IAM) (23, 38, 40–42). According to (IAM), molecular chaperones anneal the population of misfolded biopolymers by disrupting their structures and offering them multiple chances to fold. The theory based on the IAM predicts that the yield of the native state (N), after *n* iterations of folding transitions, *Y*_N_(*n*), is given by, (23, 41, 42)

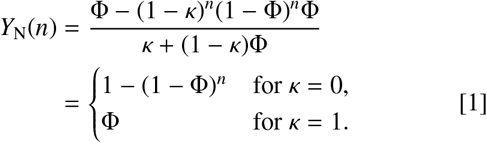

where the recognition factor, *κ*, is the ratio of chaperone-mediated transition rates associated with reversing the spontaneous transitions to the native and misfolded states (see Fig. 1), respectively. For *κ* ≈ 0, the unfolding action of chaperone is restricted to the misfolded states, the condition that holds in the case of the bacterial GroEL-GroES chaperonin system (38, 40). On the other hand, *κ* = 1 corresponds to a hypothetical chaperone that promiscuously interacts with both the misfolded and native conformations. Whereas a chaperone with *κ* = 0 increases *Y*_N_(*n*) from Φ (*n* = 1) to the unity (*n* ≫1 - a proxy for time), a chaperone with *κ* = 1 leaves the native yield unaltered (*Y*_N_(*n*) = Φ for all *n*). The latter process would correspond to a futile enzymatic cycle. For the case of CYT-19 acting on *T*. ribozyme, the value of *κ* is in the range 0 ≲ *κ* ≪ 1 (*κ* ≈ 0.1 (23)).

The generalized IAM (Eq. 1), that was mapped onto the three-state kinetic model (23), showed that although the steady state native yield of the ribozyme decreases with increasing CYT-19 concentration, the product of *the chaperone-mediated folding rate* and *the steady-state yield of native substrate* is an increasing function of chaperone concentration for both RNA and protein chaperones. This optimization condition suggests that molecular chaperones increase the production of functionally active substrates on a biologically relevant timescale, which appears to be a general principle governing the functions of a class of chaperones. Achieving the optimization comes at an expense (38). For instance, it has been argued that CYT-19-facilitated folding of ribozyme is an energetically intensive process, resulting in the hydrolysis of as many as ∼ 100 ATP molecules in order to assist the folding of a single misfolded ribozyme (37). The optimization condition, requiring substantial amount of ATP consumption, highlights the non-equilibrium nature of chaperone-facilitated dynamics of substrate molecules for a number of seemingly unrelated systems (42).

There are a plethora of studies that investigate the effects of CYT-19 on the folding of *T*. ribozyme. However, studies on the CYT-19 effect on the self-splicing of group I intron in pre-RNA is relatively scarce, except one that used a mutant ribozyme without the P5abc domain (25). The lifetime of the native state is finite, and even shortened by the helicase activity of CYT-19. Given that self-splicing of group I ribozyme occurs in a finite time, we expect that there ought to be competition between the lifetime of the native ribozyme, and the timescale for the self-splicing reaction. To our knowledge, there are no studies that explore the link between RNA chaperone mediated folding and the splicing activity. Here we fill this gap using theory. We first analyze the recent *in vitro* measurements of *T*. ribozyme folding (35–37) to extract the parameters involving chaperone-activity for use in the generalized IAM-based three-state model (23). We then use our theory to make experimentally testable predictions on the role of RNA chaperones in the processing of pre-RNA *in vivo*.

Our study shows that there are multiple competing timescales in the processing of pre-RNA. The condition for maximizing the yield of the self-spliced product depends on the lifetimes of the misfolded, native state, the rate of self-splicing reaction, and the extent of CYT-19 activity. Our theory shows that the action of molecular chaperone on RNA should be strong enough to transform the misfolded ribozyme to native state, but should not be too strong to disrupt the native state before the self-splicing of pre-RNA occurs. The theory provides a quantitatively precise condition that gives rise to the optimal level of chaperone activity to maximize the yield of spliced pre-RNA *in vivo*.

## Results

### IAM based theory using three-state kinetic model for CYT-19-assisted folding of *Tetrahymena* ribozyme

At the early stage of folding, under native conditions (determined by the solvent quality, temperature, and ion concentration) RNA molecule rapidly collapses from an unfolded state, and adopts secondary structures (Fig. 2A) (44), yielding an ensemble of intermediate states (I, represented in red (Fig. 2B)). The 3D structures that are assembled by forming tertiary contacts are divided into a non-functional ensemble of misfolded states (M, green), and the functionally competent native states (N, blue) (Fig. 2B). In the absence of RNA chaperone, the transitions from I to M (I → M with rate constant *k*_IM_) or from I to N (I → N with rate constant *k*_IN_) are effectively irreversible. The fraction of ribozymes that folds spontaneously to the N state is given by the partition factor, Φ ≡ *k*_IN_/ (*k*_IN_ + *k*_IM_). For *T*. ribozyme, Φ ≈ 0.1 (19, 45–47). Transitions between the M and N states could occur with rates *k*_MN_ and *k*_NM_, but the timescale associated with the M → N transition, is far too slow to be biologically relevant (*k*_MN_ < 0.05 min^−1^) (35, 36). Thus, the dynamic process of ribozyme folding *in vitro* is under kinetic control, which implies that a substantial portion ((1 − Φ) ∼ 0.9) of the ribozyme, starting from the I state, is trapped in the M state.

**Fig. 2.**
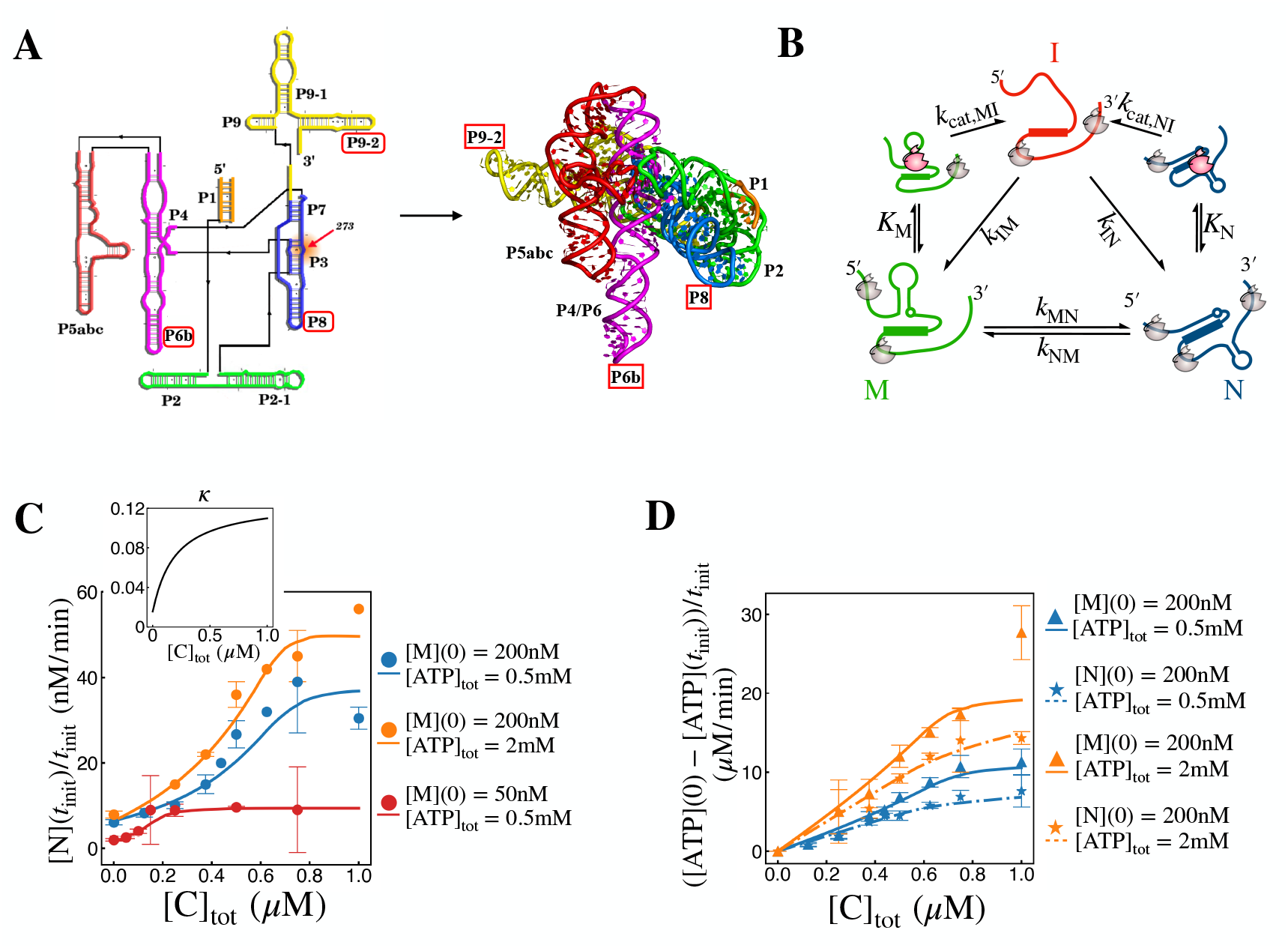
RNA structure and chaperone-assisted folding. **A**. Schematics of the folding of *T*. ribozyme. On the left is the secondary structure map and the right side represents the three dimensional native structure (PDB id: 7EZ0 (43)). **B**. A kinetic scheme for the three-state model of CYT-19-assisted folding of *T*. ribozyme used in the theoretical analysis. RNA chaperones in productive and futile binding modes are depicted in red and grey, respectively. **C, D**. Analysis of CYT-19-assisted folding of *T*. ribozyme. The curves represent the fits obtained using our model, and the symbols are the experimental data. The error bars denote two standard deviations from the mean, and the error bars are omitted for data points generated from single trials. The initial refolding rates of the ribozyme (**C**) and the ATP hydrolysis rates (**D**) measured over multiple CYT-19, ATP, and ribozyme concentrations. The initial rates of N state formation and ATP consumption were set to [N](*t*_init_)/*t*_init_ and ([ATP](0) − [ATP](*t*_init_))/*t*_init_, respectively (see SI Appendix **Model fit to data** for the definition of the early time point *t*_init_). All curves are computed with the best-fit parameters shown in Table 1. The experimental data points, obtained from (35–37), are provided in the Supplementary data.

Following the IAM-based three-state model (Fig. 2B) for the chaperone-assisted folding (23), we assume that CYT-19 binds to M and N states of the ribozyme (35) with distinct binding affinities (dissociation constants) *K*_M_ and *K*_N_. Subsequent binding of ATP to the CYT-19-ribozyme complexes, with the affinity *K*_ATP_ and the ATP-dependent helicase activity of CYT-19, revert the M and N states to the I state with rates *k*_cat,MI_ and *k*_cat,NI_. The CYT-19 and ATP concentration-dependent chaperone activity-mediated transition rates from the M and N states to the I state, represented by the *effective* rates 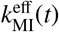 and 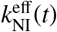, which account for the sequential events of the CYT-19 binding to ribozyme (M or N) and the catalytic activity of CYT-19 powered by ATP-consumption, are modeled using Michaelis-Menten type kinetics,

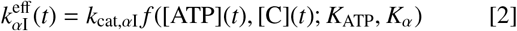

with

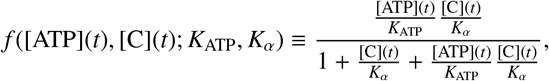

where *α* = M or N, [C](*t*) denotes the time-dependent concentrations of free CYT-19, and [ATP](*t*) is also modeled time-dependent since the ATP concentration is not constantly held in the experiments (35–37). For the derivation of Eq. 2, see SI Appendix **E**ff**ective rate constants for CYT-19 activity**, and for further detail on the calculation of [C](*t*), see SI Appendix **Mass balance of CYT-19**. The parameters *k*_cat,MI_, *k*_cat,NI_, *K*_M_, *K*_N_ and *K*_ATP_ are the catalytic rates and binding constants as shown in Fig. 2B. In the presence of chaperone activity with [C](*t*), [ATP](*t*) ≠ 0, the reactions between I, M, and N states are reversible.

Although it is not shown explicitly in Fig. 2B, CYT-19 can strongly bind to surface-exposed, protruding helices, such as P6b, P9-2, and P8 (see Fig. 2A) and unwind them by futile consumption of ATP (37). Taken together, in our theoretical framework, that implicitly takes into account the futile ATP consumption to unwind the protruding helices, ATP molecules are consumed via three pathways and obey the following rate equation,

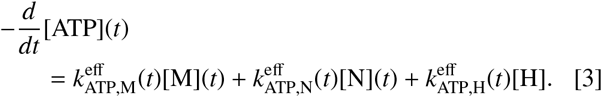

Here, 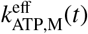 and 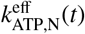, are the effective ATP turnover rates catalyzed by CYT-19 bound to the ribozyme in the M and N states, respectively, and are defined similarly to 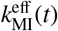 and 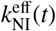 in Eq. 2 (see Eq. S7 for more precise definition). 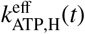 is the effective ATP turnover rate catalyzed by CYT-19 bound to the surface exposed helices. The total concentration of the surface exposed helices is given by [H] = *n*_H_[R]_tot_, where we assume that the number of secondary structure motifs, *n*_H_, is identical in the I, M, and N states. To account for the constitutively exposed helices of *T*. ribozyme, P6b, P8, and P9-2 (see Fig. 2A), we set *n*_H_ = 3 (36). It is noteworthy that Eqs. 3, S7, and S8 model the rate of ATP consumption to be proportional to the amount of CYT-19-ribozyme complex, and not to the occurrence of unfolding reactions, so that they include the contribution from unproductive futile cycles, which do not result in unfolding the ribozyme. For further detail, see the SI Appendix **Effective rate constants for CYT-19 activity**.

Finally, the kinetic and thermodynamic parameters for the model, given in Fig. 2B, are determined by simultaneously fitting the biochemical data collected from three reports (35–37) to the solution of the three-state kinetic model described by the following rate equation:

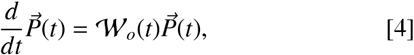

where 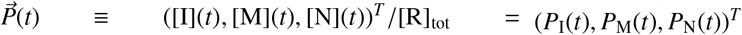 with [R]_tot_ = [I] + [M] + [N] is the vector of the normalized concentrations (or probabilities), and the rate matrix 𝓌_*o*_(*t*) is

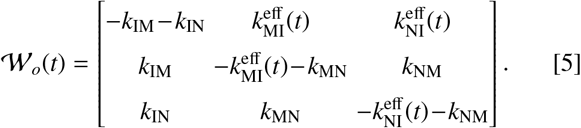

More specifically, the parameters in the theory were obtained by simultaneously fitting to 11 sets of data on the folding of *T*. ribozyme, (Fig. S1) and 2 sets of data on the folding of mutant *T*. ribozyme lacking two helices (Fig. 2C) and ATP consumption rate (Fig. 2D) with varying concentrations of CYT-19, ATP, and ribozyme, along with two constraints on the parameters: (i) Φ = *k*_IN_/(*k*_IN_ + *k*_IM_) = 0.1 (45–47); (ii) *k*_MN_/*k*_NM_ = 10 based on the observation that ∼ 90 % of the ribozyme are found in the N state at equilibrium (Fig. 2a in Ref. (36)). We extracted the best-fit parameters by minimizing the weighted sum of squared residuals (χ^2^-statistic, which is further explained in the SI Appendix **Model fit to data**). It should be emphasized that despite having about ten parameters, they were robustly obtained by simultaneously fitting to large set of data. The model captures all the experimentally observed trends (Figs. 2C,D and Fig. S1), and the best-fit parameters are consistent with experimental measurements, although some of the measurements were performed under different conditions of temperature and ion concentrations (Table. 1).

From the parameters determined from the analysis the following inferences may be made.

1. CYT-19 binds more strongly to the M state (*K*_M_ < *K*_N_), and catalyzes the unfolding reaction more rapidly (*k*_cat,NI_ < *k*_cat,MI_). The helicase activity of CYT-19 is more productive if the ribozyme is in the M state. The extracted parameters in Table 1 indicate *k*_cat,NI_/*K*_N_ < *k*_cat,MI_/*K*_M_. As a result, CYT-19 displays a greater “specificity” towards the M state with the recognition factor satisfying 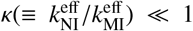. In the range of [C]_tot_ = (1 − 20) µM and [ATP]_tot_ = 0.5 mM, *κ* increases monotonically with [C]_tot_, from *κ* ≈ 0.01 to *κ* ≈ 0.1. These results are consistent with a previous study (23).
2. The binding of CYT-19 to the surface-exposed, protruding helices (P6b, P9-2, and P8. See Fig.2A) is stronger than to the ribozyme core (*K*_H_ < *K*_M_, *K*_N_), which means that the helices effectively sequester CYT-19 and hamper the CYT-19 assisted folding of the ribozyme when the amount of free CYT-19 in solution is small. The effect of protruding helices manifests itself as a sigmoidal increase in the folding rate with increasing CYT-19 concentration (Fig. 2C). Note that the initial rates of ATP hydrolysis in the presence of M and N states are comparable. This is because a large portion of the ATP turnover rates are contributed by CYT-19 bound to the protruding helices (*K*_*H*_ ≪ *K*_*M*_ ≪ *K*_*N*_. See Table 1), especially when the [C]_tot_ concentration is low (Fig. 2D).

Our model, in which the number of exposed helices per mutant ribozyme devoid of P9-2 and P6b is set to unity (*n*_H_ = 1), qualitatively reproduces the experimentally observed suppression of the refolding rate of wild type ribozyme in comparison with the mutant (Fig. S1E), and the lower ATP consumption rate in the mutant (Fig. S1F). We find that ATP catalysis occurs five times faster in the productive CYT-19 binding to the ribozyme than in the binding to the protruding helices (*k*_cat,P_ ≈ 5*k*_cat,H_).

**Table 1.** The best-fit parameters extracted by fitting the experimental data at 25°C with 1 mM Mg^2+^ and 2 mM Mg^2+^ in Ref. (35) and with 2 mM Mg^2+^ in Refs. (36, 37).

| | $k_{\text{cat},MI}$<br>($\text{min}^{-1}$) | $k_{\text{cat},NI}$<br>($\text{min}^{-1}$) | $K_H$<br>(nM) | $K_M$<br>(nM) | $K_N$<br>(nM) | $*k_{NM}$<br>($\text{min}^{-1}$) |
| --- | --- | --- | --- | --- | --- | --- |
| Best-fit | 11 | 1.4 | 7.3 | 50 | 5000 | $3.6 \times 10^{-3}$ |
| | $*k_{IM}$<br>($\text{min}^{-1}$) | $*k_{IN}$<br>($\text{min}^{-1}$) | $*k_{MN}$<br>($\text{min}^{-1}$) | $K_{ATP}$<br>($\mu\text{M}$) | $k_{\text{cat},H}$<br>($\text{min}^{-1}$) | $k_{\text{cat},P}$<br>($\text{min}^{-1}$) |
| Best-fit | 4.7 | 0.53 | 0.037 | 900 | 27 | 140 |
| exp. | $^{a}1.5 \pm 0.3$ | $^{a}0.37 \pm 0.10$ | $^{b}0.05$ | $^{c}50 - 500$ | $^{c}6 - 600$ | $^{c}6 - 600$ |

\* In the fitting procedure, we fix the ratios $k_{MN}/k_{NM} = 10$ and $\Phi = k_{IN}/(k_{IN} + k_{IM}) = 0.1$ . <sup>a</sup> Ref. (45), with 10 mM $\text{Mg}^{2+}$ . <sup>b</sup> Ref. (35), with 1 mM $\text{Mg}^{2+}$ . <sup>c</sup> Refs. (48, 49). The relations among the best-fit parameters, $k_{\text{cat},NI} < k_{\text{cat},MI}$ , $K_H < K_M < K_N$ , and $k_{ATP,H}^{\text{eff}} < k_{ATP,P}^{\text{eff}}$ are also confirmed in the distribution of good-fit parameters acquired from the exhaustive sampling using Markov Chain Monte Carlo (MCMC) method (Fig. S2).

### Predictions for CYT-19-facilitated folding and self-splicing of pre-RNA

The aforementioned experiments (35–37) were concerned with the folding of *T*. group I intron ribozyme (35–37), which catalyzes its own excision from the precursor RNA (pre-RNA) (51). Here, we extend the IAM-based model to predict the effect of CYT-19 activity on the self-splicing of pre-RNA.

To study the effect of CYT-19 on pre-RNA, we refer to the model suggested by Pan *et al*. who performed *in vitro* folding measurements (19). The model assumes that *T*. ribozyme containing pre-RNA undergoes transitions among the ensembles of intermediate states (I), two different misfolded states (M_1_ and M_2_), and the native state (N) (Fig. 3A). In addition, CYT-19 is assumed to bind to the M_1_, M_2_, and N states with dissociation constants 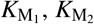 and *K*_N_, respectively. Subsequent CYT-19-mediated transition rates from the M_1_, M_2_ and N states to the I state are denoted by the effective rate constants 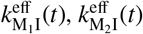 and 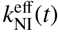 which are given in the form,

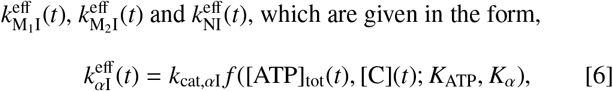

with *α* = M_1_, M_2_ or N. In the absence of experimental data, we set the concentration of the protruding helices to 0 in the calculation of [C](*t*) ([H]_tot_ = 0. See Eq. S11 in SI Appendix **Mass balance of CYT-19** for further detail).

**Fig. 3.**
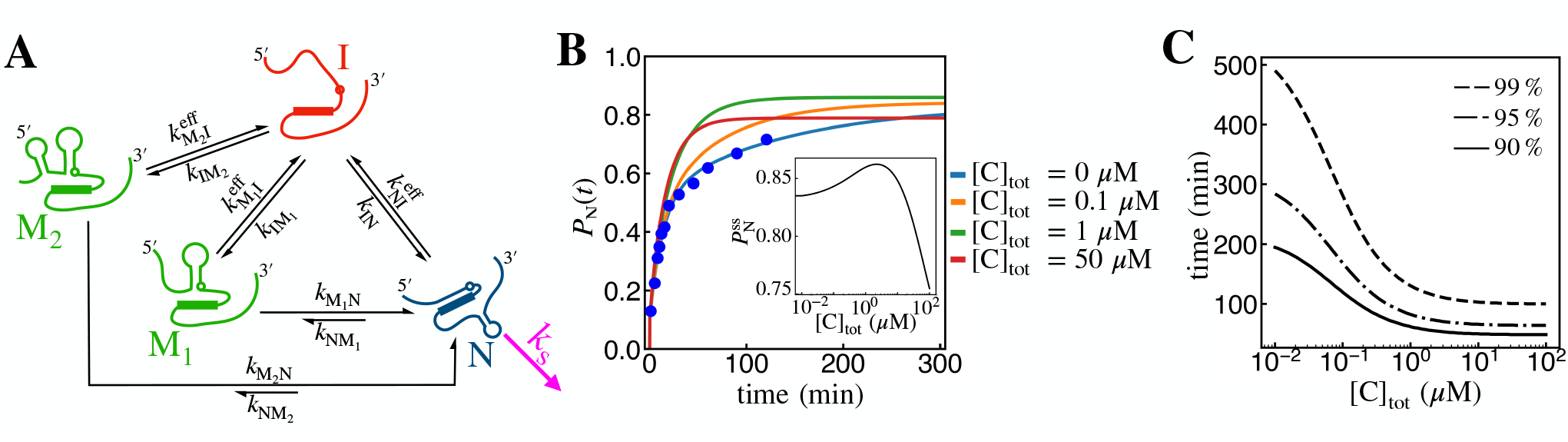
CYT-19-assisted folding and self-splicing of pre-RNA. **A**. Schematic network model for chaperone-assisted folding coupled to self-splicing of pre-RNA. The self-splicing reaction can occur at the splice site (open circle), and could be inhibited partly by the hairpin structure formed between the 5’ exon and the internal guide sequence (filled box) (50). **B**. Fraction of pre-RNA in the N state, *P*_N_, as a function of time, for varying concentrations of CYT-19. The inset in **B** shows the steady state yield of native pre-RNA, *P*^ss^, as a function of [C]_tot_. **C**. The first passage time *t*_fpt_ that satisfies, **the probability of spliced state**, *P*_SP_(*t*_fpt_) = 0.90, 0.95 and 0.99 are shown as a function of [C]_tot_. The values of *t*_fpt_ decrease sharply as [C]_tot_ increases. All the curves are plotted using Eq. 7 with [ATP]_tot_ = 2 mM and [R]_tot_ = 200 nM, and the kinetic parameters are as described in Table 2. The data points in **B** (blue filled circles) are from Ref. (19).

We consider the experimental setup in which [R]_tot_ of pre-RNA is incubated with [C]_tot_ of CYT-19 and [ATP]_tot_ of ATP. The time evolution of the pre-RNA concentrations in the four states, [I](*t*), [M_1_](*t*), [M_2_](*t*) and [N](*t*), are described by

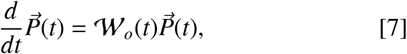

where 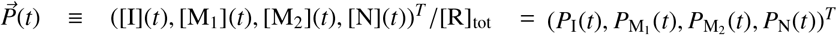 is the vector of the normalized concentrations, and the transition matrix, 𝓌_*o*_(*t*) is given by,

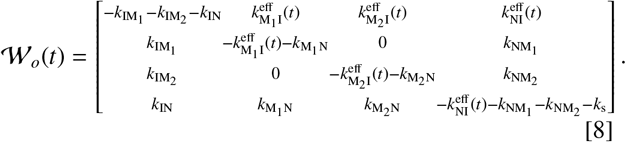

The splicing rate is set to *k*_s_ = 0 min^−1^ when the self-splicing reaction is blocked, e.g., by omitting the required guanosine cofactor. But, in the presence of guanosine co-factor, the splicing rate is set to *k*_s_ = 2.5 min^−1^.

The filled circles in blue in Fig. 3B are the time evolution data of native pre-RNA, *P*_N_(*t*), reported by Pan *et al*. (19) who measured *P*_N_(*t*) starting from the initial condition *P*_I_(0) = 1, while blocking the self-splicing reaction. It was shown, using theoretical arguments and experimental data, that a small fraction of pre-RNA (Φ ≈ 0.1) must fold rapidly to the N state with *k*_IN_ = 60 min^−1^, in accord with theoretical estimates (44, 52), and observed subsequently in a single-molecule experiment (31). The remaining fraction transition slowly into the N state in a bi-phasic manner (19), suggestive of two distinct ensembles of misfolded states (M_1_ and M_2_). In the absence of RNA chaperone, the pre-RNA in the M_1_ and M_2_ states transition slowly to the N state with rates 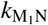 and 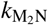. The values for the associated rate constants listed in Table 2 are obtained by fitting the measured time trace of the fraction of the native state to the solution of Eq. 7. Specifically, we minimize the sum of the squared residuals between the measurements and predicted curves, with the constraints *k*_IN_ = 60 min^−1^, 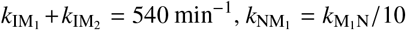 and 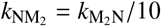.

**Table 2.** Parameters for folding and self-splicing of pre-RNA in the presence of CYT-19.

| | $k_{M_1N}$ | $k_{M_2N}$ | $k_{NM_1}$ | $k_{NM_2}$ | $k_{IM_1}$ | $k_{IM_2}$ | $k_{IN}$ |
| --- | --- | --- | --- | --- | --- | --- | --- |
|  | (min<sup>-1</sup>) | (min<sup>-1</sup>) | (min<sup>-1</sup>) | (min<sup>-1</sup>) | (min<sup>-1</sup>) | (min<sup>-1</sup>) | (min<sup>-1</sup>) |
| Best-fit | 0.07 | 0.007 | 0.007 | 0.0007 | 264 | 276 | 60 |

| | $k_{cat,M_1I}$ | $k_{cat,M_2I}$ | $k_{cat,NI}$ | $K_{ATP}$ | $K_{M_1}$ | $K_{M_2}$ | $K_N$ |
| --- | --- | --- | --- | --- | --- | --- | --- |
|  | (min<sup>-1</sup>) | (min<sup>-1</sup>) | (min<sup>-1</sup>) | (μM) | (μM) | (μM) | (μM) |
|  | 11/S | 11/S | 1.4/S | 900 | 50S | 50S | 5000S |

Since the folding of pre-RNA was measured at $[Mg^{2+}] = 6$ mM (19), the turnover rates and binding affinities associated with CYT-19 activity are adjusted using the factor $S = e^{m([Mg^{2+}] - 2mM)}$ with $m \approx 1$ mM<sup>-1</sup> (53), such that $k_{cat,M_1I}$ , $k_{cat,M_2I}$ , $k_{cat,NI}$ , $K_{M_1}$ , $K_{M_2}$ , are scaled to $K_N \rightarrow k_{cat,M_1I}/S$ , $k_{cat,M_2I}/S$ , $k_{cat,NI}/S$ , $K_{M_1}S$ , $K_{M_2}S$ , $K_NS$ .

For the model shown in Fig. 3A, we assume that CYT-19, when present, binds to the M_1_, M_2_ and N states and induce unfolding transitions. For the parameters associated with CYT-19 dynamics, we used the values from Table 1, assuming that CYT-19 associated parameters for M_1_ and M_2_ are identical (for further detail, see Eq. 6 and Table 2). Increase in the CYT-19 activity facilitates the production of the N state at early times, but this trend does not continue to hold at later times (Fig. 3B). As a consequence, the steady state yield of the native state 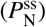 depends non-monotonically on [C]_tot_. At low [C]_tot_, when CYT-19 predominantly binds to the M state, 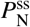 increases with [C]_tot_; however, at high [C]_tot_, when substantial CYT-19-mediated unfolding of the N state ribozyme takes place, 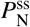 decreases with [C]_tot_ (inset in Fig. 3B). This key prediction suggesting an optimal value of CYT-19 for folding of the ribozyme is an emergent property of our theory.

In the presence of the guanosine co-factor, pre-RNA in the N state self-splices irreversibly with the rate *k*_s_ ≈ 2.5 min^−1^ (54). The probability that pre-RNA has undergone self-splicing can be calculated using 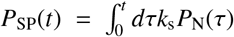. Since the self-splicing reaction is irreversible and the system is closed, *P*_SP_(*t* ≫ 1) reaches 1 in the steady states, for all CYT-19 concentrations. It is important to note that unlike 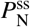 that displays non-monotonic dependence on [C]_tot_ (inset of Fig. 3B), *P*_SP_(*t*) increases monotonically. The first passage time, *t*_fpt_, that satisfies the conditions, *P*_SP_(*t*_fpt_) = 0.90, 0.95, and 0.99, decreases monotonically with [C]_tot_ (Fig. 3C). This is because most of the pre-RNA in the N state would undergo self-splicing before being unfolded by CYT-19, with *k*_s_ = 2.5 min^−1^ ≫ *k*_cat,NI_ ≈ 0.02 min^−1^. These results show that functional requirements override the kinetic criterion for folding. Hence, the ribozyme has evolved to optimize the rate of the splicing reaction.

## Discussion

We generalized the iterative annealing mechanism to characterize the chaperone-assisted folding of RNA and the associated consumption of ATP by chaperone activity. Our theory is consistent with data on the folding and splicing of ribozymes (Fig. 2 and Fig. 3), and highlights the competition between the chaperone-mediated unfolding and self-splicing reactions in the native state (Fig. 3). In the following sections, we further explore the important implications of our theory, including the thermodynamic cost of CYT-19 activity, and the effect of chaperone activity in RNA metabolism *in vivo*, where the yield of the spliced product is determined by the self-splicing, folding, degradation, and chaperone-mediated unfolding reactions.

### Thermodynamic cost of CYT-19 activity needed to fold a single misfolded ribozyme

Chaperone activity generated by ATP hydrolysis, which reshapes the distribution of RNA conformations in the rugged folding landscapes, is a non-equilibrium phenomenon (23, 35, 55). A quantity of great relevance, which goes to the heart of the quantities that are optimized *in vivo*, is the thermodynamic cost of correcting the errors due to the formation of non-native interactions in a single misfolded ribozyme to the native state, which we quantify using the average number of ATPs consumed by CYT-19, ⟨*A*_M→N_⟩. By calculating the thermodynamic costs precisely, it is possible to quantitatively assess how cells balance the energy costs to fold ribozymes in order to execute the self-splicing reaction.

Since the spontaneous transitions from N to I or M are rare, we set 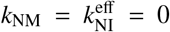 in Eq. 4. Counting only the ATPs consumed by the productively bound CYT-19, and setting the rate of ATP processing in the M state by 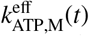 (Eq. S7), we calculated the average number of ATP consumed in converting M → N, ⟨*A*_M→N_⟩ as follows (See Eq. S17 in the SI Appendix, **Mean first passage time** for the derivation),

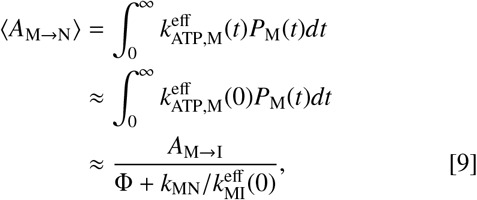

where 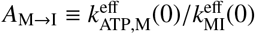 corresponds to the number of ATPs catalyzed in a single event of chaperone-mediated unfolding (M → I). To obtain the simplified expression in the last line of Eq. 9, we assume that chaperone-mediated unfolding rates only vary weakly with time, which implies that 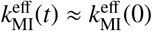 and 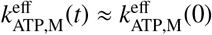. These approximations are validated by the similarity between the two ⟨*A*_M→N_⟩s (solid and dashed lines in Fig. 4A,B) calculated based on the expressions in the first and the last lines of Eq. 9.

**Fig. 4.**
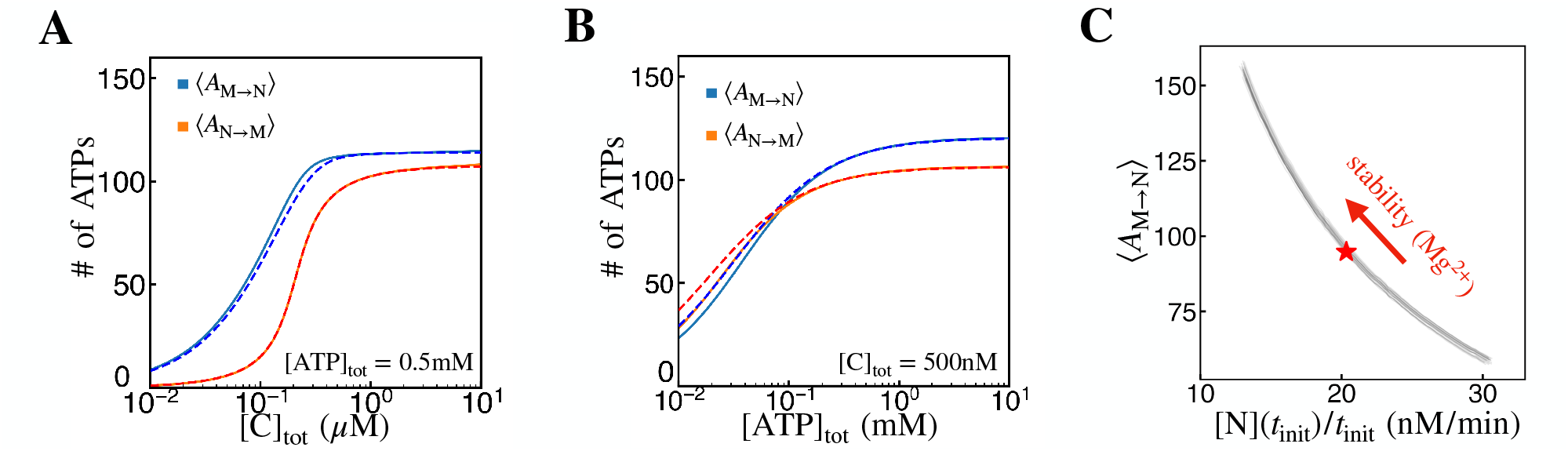
ATP consumption during the CYT-19-assisted folding of ribozyme. **A, B**. The average number of ATP molecules required to transform a single misfolded ribozyme to the native state (⟨*A*_M→N_⟩), or a single native ribozyme to the misfolded state (⟨*A*_N→M_⟩), as functions of (A) CYT-19 and (B) ATP concentrations (solid lines). The dashed lines in (A) and (B) are the approximate expressions in Eqs. 9 and 10. **C**. Plot of *A* versus the initial rate of N state formation ([N](*t*)/*t*), as functions of ribozyme stability. Ribozyme stability was modulated by the factor 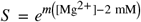, with *m* = 1 mM^−1^ and [Mg^2+^] ranging from 1.5 to 2.5 mM. The star denotes the condition of [Mg^2+^] = 2 mM (*S* = 1). The individual lines represent 20 of the parameter vectors sampled using MCMC, 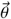, with the constraint 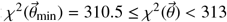, where 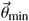 is the best-fit parameter vector shown in Table 1. For details on MCMC sampling, see the SI Appendix **MCMC sampling**.

With the best-fit parameters listed in Table 2, we compute ⟨*A*_M→N_⟩ for varying concentrations of CYT-19 and ATP (Fig. 4A, B). ⟨*A*_M→N_⟩ increases with CYT-19 and ATP concentrations, and saturates to a constant value *A*_M→I_/Φ ≈ 125, where *A*_M→I_ ≈ 12.5 and Φ = 0.1. Our prediction is in good agreement with the experimental estimate (37). Thus, RNA chaperone consumes ATP lavishly in order to ensure that the ribozyme folds, which is not unlike the bacterial chaperonin, GroEL (23).

If the rate of transition M → N is negligible 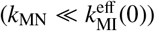, the number of ATPs required to refold the misfolded ribozyme to the native state is, ⟨*A*_M→N_⟩ ≈ *A*_M→I_/Φ ≈ 125 (Eq. 9, Fig. 4). In comparison, for RuBisCO with Φ = 0.02 − 0.05 (38),

M → I transition demands consumption of 3 – 4 ATPs per cycle (*A*_M→I_ ≈ 3 − 4) (56) for the case of the bacterial GroES-GroEL system (57). Thus, we estimate that about 60 – 200 ATPs are required for GroEL to anneal a single misfolded RuBisCO to its native form. It was argued elsewhere (38) that the large number of ATP needed to fold RuBisCo (or other proteins) is a small price to pay compared to the free energy cost needed to synthesize the protein in the first place. Similar reasoning holds for RNA synthesis as well.

With the assumption that *k*_MN_ = *k*_cat.MI_ = 0, we also obtain an expression for ATP consumption involving the chaperone-mediated transition of N → M,

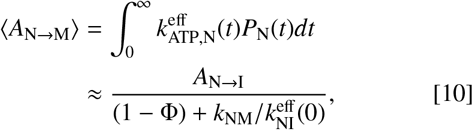

with 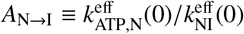.

The large amount of ATP consumption by chaperones (Eqs. 9, 10, and Fig. 4) raises a couple of points that are worthy of further discussion:

1. *A*_N→I_(≈ 97) ≫ *A*_M→I_(≈ 12.5) shows that more amount of ATPs consumed for CYT-19 to disrupt the ribozyme in the N state than if the ribozyme is misfolded, which accords well with their stabilities. We also surmise that futile consumption of ATP must be prevalent in the helicase activity of CYT-19, which involves disruption of the stable secondary or tertiary motifs of ribozyme. The prediction that there are futile cycles is common to many molecular machines (42). It is also noteworthy that the ratio *A*_N→I_/*A*_M→I_ is similar to that between the concentrations of ribozyme in the N and M states at equilibrium, [N]_eq_/[M]_eq_ ≈ 9 (36).
2. Since *k*_MN_, *k*_NM_ ≪ 1, Eqs. 9 and 10 may be simplified as ⟨*A*_M→N_⟩ ≈ *A*_M→I_/Φ and ⟨*A*_N→M_⟩ ≈ *A*_N→I_/(1 − Φ). From Figs. 4A and 4B, it follows that ⟨*A*_M→N_⟩ ≈ ⟨*A*_N→M_⟩. This shows that, although the cost of unfolding a ribozyme in the M state is smaller than unfolding a ribozyme in the N state, there are more ribozymes in the M state than in the N state (Φ ≪1). As a result, the average number of ATPs required by a chaperone to unfold a substrate molecule in the M and N states are comparable.

The more stable and compact an RNA is, the weaker should be chaperone binding to the ribozyme, and less frequent would be the transitions between I, M, and N states. In experiments, RNA stability is controlled by changing the Mg^2+^ concentration ([Mg^2+^]). Although *k*_ATP,P_ and *K*_ATP_ do not change, Mg^2+^ affects the transition rates of RNA dynamics and binding affinities of CYT-19. The factor 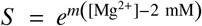, where the m-value (*m* ≈ 1 mM^−1^) quantifies the effect of Mg^2+^ on the stability of ribozyme in the folded states (53), may be used to calibrate the transition rates and affinities of CYT-19 to ribozyme at Mg^2+^ concentration other than at the physiological concentration of Mg^2+^, [Mg^2+^] = 2 mM. With increasing [Mg^2+^], ⟨*A*_M→N_⟩ increases, and the production rate of N state measured at an early stage ([N](*t*_init_)/*t*_init_) is reduced, which implies that the thermodynamic cost of creating RNA with enhanced stability is greater, resulting in the overall slowing down of the overall process (Fig. 4C). Our finding, summarized in Fig. 4C, is in good agreement with the trend observed in the experiment (Fig. 4A in Ref. (37)).

### IAM-based model for self-splicing of pre-RNA *in vivo*

Upon folding to their respective native ribozyme structures, the introns undergo self-splicing reactions (25, 26, 58). If the folding and self-splicing processes are too slow, RNA can be removed either by degradation or dilution from cell growth (59, 60). In the following, we consider the effect of chaperone on the self-splicing ‘yield’ of pre-RNA *in vivo*, which hinges on the competition between spontaneous folding, chaperone-mediated unfolding, degradation, and self-splicing reactions.

In our minimal model, upon transcription pre-RNA rapidly reaches the I state, which is the entry point in the kinetic network that describes coupled folding and splicing (Fig. 3A). We assume that the pre-RNA transitions between the states I, M, N, and SP at the rates that were extracted through the analysis of the *in vitro* model, and is degraded at a rate *k*_d_ from all the four states (Fig. 5A). The pre-RNA molecules evolve with time as,

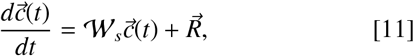

where 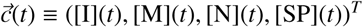,

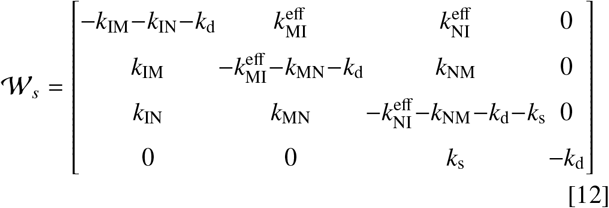

and 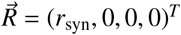 is the input source current to the I state (see the cyan on the left in Fig. 5A) that represents the rate of pre-RNA being supplied upon transcription. The steady state concentrations [I]^ss^, [M]^ss^, [N]^ss^, [SP]^ss^ are obtained from 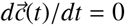 in Eq. 11. The total concentration 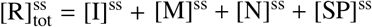 can be equated as 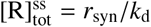.

**Fig. 5.**
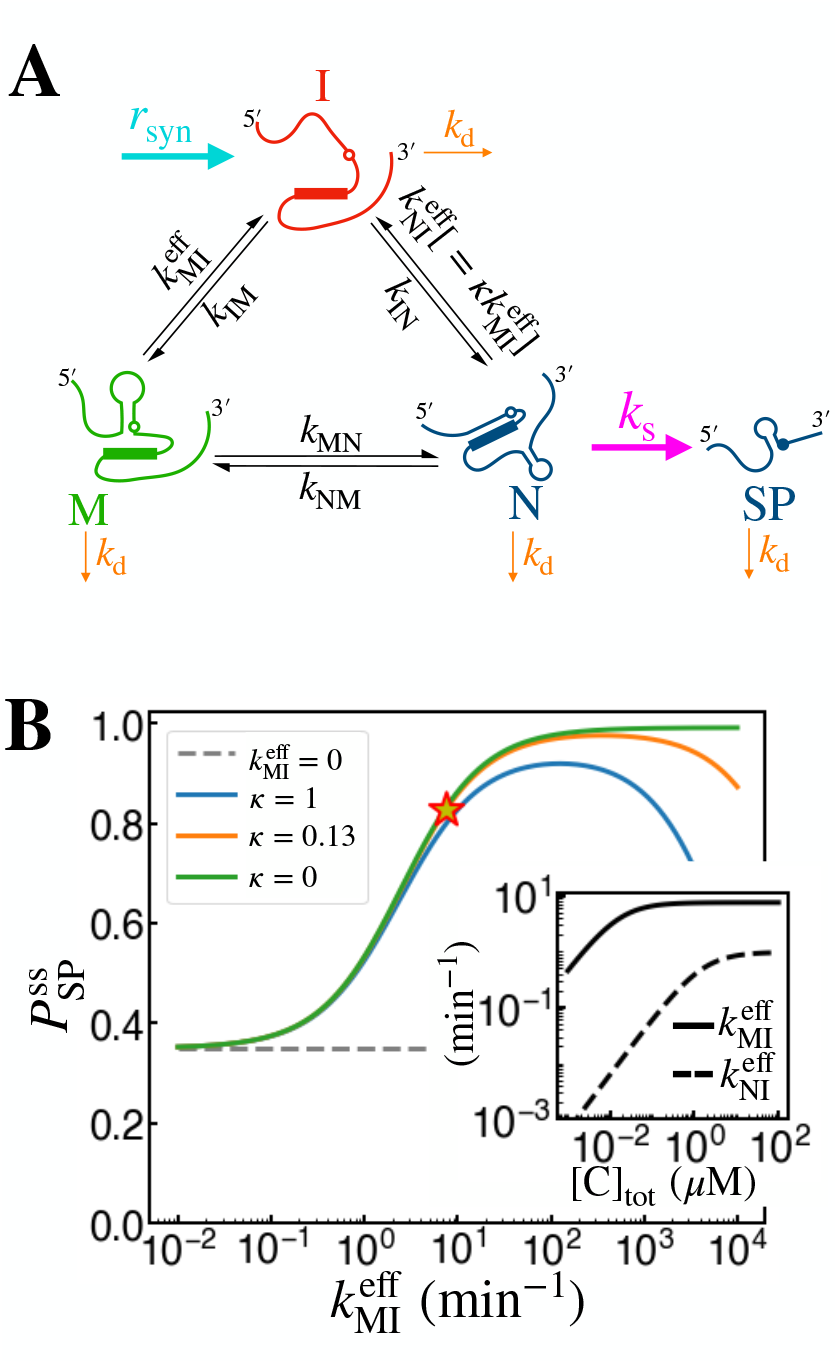
Self-splicing *in vivo*. **A**. Schematic for chaperone-mediated folding and self-splicing of pre-RNA *in vivo*. **B**. Self-splicing yield 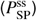 as a function of chaperone activity 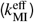, at different values of the recognition factor *κ*. The parameters other than 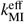 and 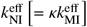 are set to *k*_IN_ = 60 min^−1^, *k*_IM_ = 540 min^−1^, *k*_MN_ = 0.07 min^−1^, *k*_NM_ = 0.007 min^−1^, *k*_d_ = 0.18 min^−1^, and *k*_s_ = 30 min^−1^. The inset in **B** shows the effective unfolding rates computed using Eq. 2, where we assumed [C] = [C]_tot_ and [ATP] = 2 mM, and used the best-fit parameters in Table 1.

Experimental measurements (25, 59, 60) show that the self-splicing rate is greater than the folding (I → M or I → N) rates, and that folding occurs more rapidly than the degradation (*k*_d_ < *k*_IN_, *k*_IM_ < *k*_s_). The chaperone activity is quantified by the effective rate constants, 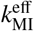 and 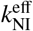, with implicit dependence on the ATP and chaperone concentrations. Here we set the splicing rate to *k*_s_ = 30 min^−1^ that was measured in the cellular environment (61). With *k*_d_ = 0.18 min^−1^ (60), and the folding rates of pre-RNA, *k*_IN_, *k*_IM_, *k*_MN_ and *k*_NM_, obtained from the fit to *in vitro* data (Fig. 3B), we calculate 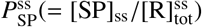 for *Tetrahymena* pre-RNA as a function of 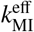 (Fig. 5B).

Our calculation shows that the chaperone activity can increase the steady state yield of the spliced ribozyme, 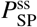 up to a certain level by facilitating ribozyme folding. However, when the chaperone activity exceeds a critical value, it leads to unfolding of the native ribozyme, and reduces the 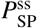, especially when *κ* > 0, which engenders a non-monotonic variation of *P*^ss^ when the chaperone activity increases (Fig. 5B).

Under the condition of *k*_MN_, *k*_NM_ ≪1 and *k*_d_ ≪ *k*_IM_, *k*_s_, the complicated expression for 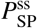 simplifies greatly. For the full expression of 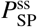 and the derivation of Eq. S23, see Eqs. S21 and S23 in the SI Appendix **Self-splicing yield**. It can be shown that 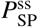 is indeed a non-monotonic function of chaperone activity 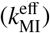, which is maximized at

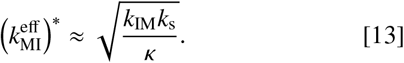

Eq. S23 can be rearranged as 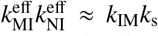, which offers greater insights into the conditions for the chaperone to maximize 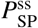. On the one hand, the chaperone-induced unfolding rate of the misfolded state should be sufficiently large compared to the rate of folding 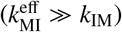, so that the pre-RNA can escape from the misfolded state. On the other hand, the unfolding rate of the native state 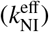 should not overwhelm the rate of self-splicing (*k*_s_), which would cause the pre-RNA in the N state to unfold before splicing. The interplay between these opposing factors determines the optimal condition for function.

More generally, we can consider a pre-RNA under weakly stabilizing conditions, in which spontaneous unfolding from N or M to I state is significant and chaperones are not required. Even in this case, our formulation based on the kinetic scheme in Fig. 5A and the optimal chaperone activity summarized in Eq. S23 are applicable by considering that 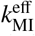 and 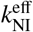 represent the sum of the spontaneous unfolding and chaperone-mediated unfolding rates of the pre-RNA ribozyme. Therefore, we surmise that ‘moderate’ activity of CYT-19 is required to maximize the self-splicing yield of the pre-RNA, which is the condition that ought to be met for other downstream processes.

Although the value of 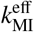 is varied over a broad range in Fig. 5B, 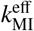 is upper-bounded to *k*_cat,MI_ (Eq. 2). At [ATP] = 2 mM, it saturates to 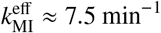 (Fig. 5B inset). Along with 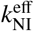, we find that 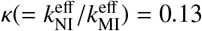 at the saturating CYT-19 concentration. Notably, the self-splicing yield calculated from 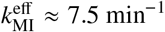 with *κ* = 0.13 is 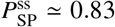 (star symbol in Fig. 5B), which is close to the maximum, 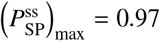 (the maximum point of the orange line in Fig.5B). The discrepancy between the two numbers, albeit small, requires some discussion. It may simply reflect that *k*_cat,MI_, the speed limit of 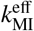, is smaller than 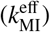 for the case of CYT-19 acting on group I intron of pre-RNA. Otherwise, given the promiscuity of CTY-19 action on surface-exposed helices of structured RNAs, CYT-19 activity cannot be fully optimized for all the structured RNAs in terms of 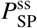. More specifically, for RNAs that fold correctly on their own with Φ ∼ 1, which is tantamount to k_IM_ ∼ 0, the optimal chaperone activity, 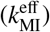* (Eq. S23), approaches 0. In this case, CYT-19 activity (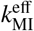 increased beyond 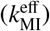*) will lead to a reduction in the self-splicing yield while catalyzing the futile cycles of ATP hydrolysis. Thus, it may be argued that the extent of CYT-19 activity (star symbol in Fig. 5B) reflects the balance between facilitating the folding of RNA with Φ close to 0, and limiting the futile unfolding reactions on RNA with Φ close to 1.

Our theory that incorporates the splicing reaction in the folding network clarifies the effect of the recognition factor *κ* on the self-splicing yield 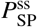. We find that the differential effect of *κ* on 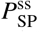 manifests itself only at *hypothetical* values of 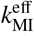 that exceed the turnover rate of CYT-19, *k*_cat,MI_. An RNA chaper one with a smaller *κ* engenders a larger 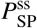 for *Tetrahymena* pre-RNA if such a chaperone exists in nature (Fig. 5B, and see SI Appendix **Self-splicing yield**). Alternatively, for a given chaperone, a ribozyme with smaller *k*_*s*_ and *k*_IM_ could reduce the value of 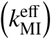* (Eq. S23), rendering the differential effect of *κ* more pronounced. It is noteworthy that the product between the chaperone-mediated folding (relaxation) rate of RNA molecule (λ) and the steady state yield of the native state 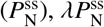, which was previously highlighted as the key quantity for optimization in the chaperone-facilitated folding in principle depends on *κ* in a self-consistent manner. It satisfies 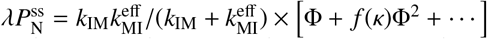 (see Eq. S28 and Fig. S3); however, when Φ is small, the dependence of 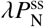 on *κ* is in effect negligible (see SI Appendix **Dependence of** 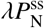 **on** *κ* for more details). *In vitro* measurements are consistent with Φ ≈ 0.1, and hence the Φ^2^ term is, for all practical purposes, negligible. For stringent protein substrates, Φ is, on the order of 0.02 (38), which even more justifies the neglect of Φ^2^ for protein chaperones. Hence, we posit that both RNA and protein chaperones evolved to maximize 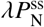.

Finally, the effect of exon sequences on the folding kinetics of pre-RNA can potentially be significant (18, 62, 63), but was not considered in this study. However, their effects may be included by suitably modifying the appropriate rates. Nevertheless, our analyses and predictions, presented in Figs. 3, 4 and 5 are amenable to experimental test. The IAM-based framework presented here illuminates the interplay between chaperone-facilitated dynamics of ribozyme and self-splicing, and could be further extended to analyze experimental data in more complex network models, which are likely required in cases involving interplay between substrate folding, multiple chaperones, and function.

## Summary

We developed a theory, based on kinetic network models that couples folding and catalysis, to determine the interplay between chaperone-mediated folding of pre-RNA and the downstream self-splicing reaction. We discovered a tension between chaperone activity and the functional requirement for sufficient yield of the self-spliced RNA. The chaperone activity must be large enough to unfold the kinetically trapped structure but not be too large to disrupt the native state because the latter is needed for the self-splicing reaction. Because the unwinding propensities of the misfolded and folded states of the pre-RNA are determined by their (kinetic) stabilities, it follows that the life time of the native state must be greater than the time needed for the splicing reaction to go to completion. The theory, which elucidates these ideas quantitatively, shows that RNA chaperones have evolved to satisfy the dual requirement of converting the misfolded structures to functionally competent folded states without disrupting the native state structures.

## Acknowledgments

This work was supported by the KIAS individual Grants CG067102 (Y.S.) and CG035003 (C.H.) at Korea Institute for Advanced Study. DT thanks the National Science Foundation (CHE-1900033) and the Welch Foundation through the Collie-Welch Chair (F-0019) for supporting this research. We thank the Center for Advanced Computation in KIAS for providing computing resources.

## SI Appendix

### 1. Effective rate constants for CYT-19 activity

We derive an expression of the Michaelis-Menten type for the effective rate constant, 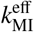, that quantifies the CYT-19-mediated unfolding of the misfolded (M state) ribozyme. The unfolding reaction can be broken down into a series of molecular events: (i) the binding of CYT-19 to the ribozyme, (ii) the binding of ATP to the catalytic site of CYT-19, and (iii) the unfolding of the M state ribozyme catalyzed by CYT-19. With 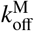 and 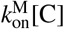 denoting the unbinding and binding rates of CYT-19 to the M state ribozyme, and 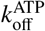 and 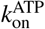 [ATP] denoting the unbinding and binding rates of ATP to the CYT-19-ribozyme complex, the reaction scheme of CYT-19-mediated unfolding reads,

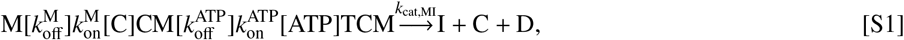

where T and D denote ATP and ADP, and CM and TCM are the CYT-19-ribozyme and ATP-CYT-19-ribozyme complexes, respectively. Based on the structural and biochemical studies, which suggested that the CYT-19 catalyzed unwinding of RNA is limited to short RNA duplexes, we assumed that the CYT-19 activity is non-processive (64, 65).

If the ribozyme molecules are initially in the M state (i.e. [M](*t* = 0) = [R]_tot_), and the concentrations [C] and [ATP] remain effectively constant in time, the time evolution of the probability vector, 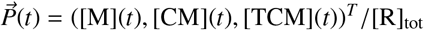 is described by the differential equation with an initial condition 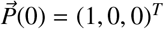,

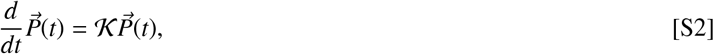

where

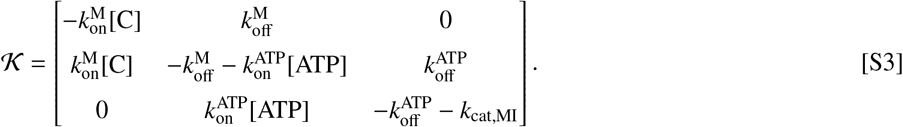

The accumulation of the I state can be quantified using 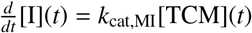, and the mean first passage time for the RNA molecule to reach the I state is given by 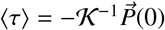 (see the SI Appendix **Mean first passage time**). The effective rate of CYT-19-mediated unfolding of M state ribozyme is then given by 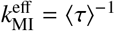.

Provided that the binding and unbinding of ATP, CYT-19, and the ribozyme are pre-equilibrated prior to the final catalytic reaction (i.e. *k*_cat,MI_ ≪ 1), we obtain the expression

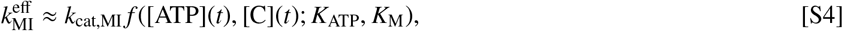

with

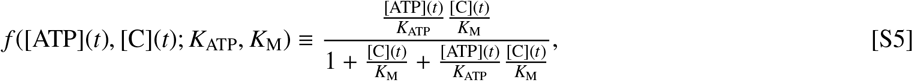

where 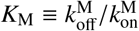 is the dissociation constant of CYT-19 from the M state ribozyme, and 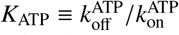 is the dissociation constant of ATP from the CYT-19-ribozyme complex. Here, the binding and unbinding of the chaperone to RNA and ATP to chaperone are assumed to be pre-equilibrated relative to the time scale of the other dynamics (66).

We note that the limit of slow unbinding of CYT-19 from the ribozyme (i.e. 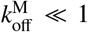) leads to the expression used by Chakrabarti *et. al*., (23)

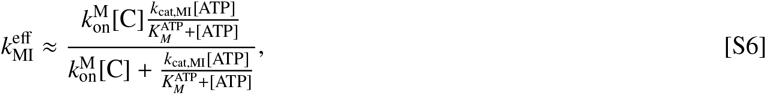

where 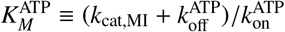. The effective rates of unfolding of native ribozyme 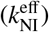 and ATP hydrolysis 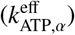 are obtained based on the same argument used in deriving 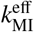.

Here, we the provide expressions for the effective ATP turnover rates, 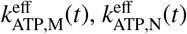, and 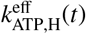, in Eq. 4 of the main text. 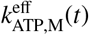 and 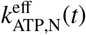 are, respectively, the effective ATP turnover rates catalyzed by CYT-19 bound productively to the and N states, and are given as

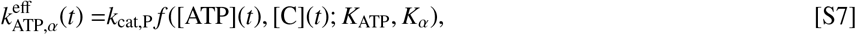

with *α* = M or N, where *k*_cat,P_ denotes the ATP hydrolysis rate of CYT-19 bound productively to the ribozyme. CYT-19 bound to the exposed helices consumes ATP with the effective rate constant 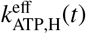

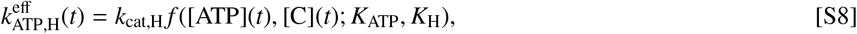

where *k*_cat,H_ is the ATP hydrolysis rate of the exposed helix-bound CYT-19. The dissociation constant of CYT-19 from the helices is given by *K*_H_. The mathematical form of *f* ([ATP](*t*), [C](*t*); *K*_ATP_, *K*_*α*_) with *α* = M, N, and H is given in Eq. S5, except that the arguments are different.

### 2. Mass balance of CYT-19

Unlike the total chaperone concentration, [C]_tot_, that stays constant in time, the concentration of free chaperone, [C](t), varies with time. Specifically, we calculate [C](t) at each time point *t* by the mass balance relation,

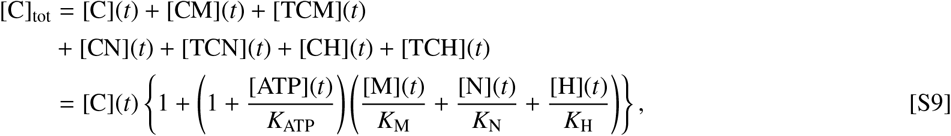

where [CN](*t*) and [CH](*t*) are the concentrations of CYT-19 complexed with the N state and the exposed helices, respectively. [TCN](*t*) and [TCH](*t*) are the concentrations of complexes where ATP molecules are bound. [M](*t*), [N](*t*), and [H](*t*) denote the concentrations of the *free* ribozymes in M and N states and *free* exposed helices without being complexed to chaperones. *K*_N_(= [C][N]/[CN]), *K*_M_(= [C][M]/[CM]), and *K*_H_(= [C][H]/[CH]) denote the respective dissociation constants from CYT-19. By expressing the free ribozyme concentration in each state in terms of [C](*t*),

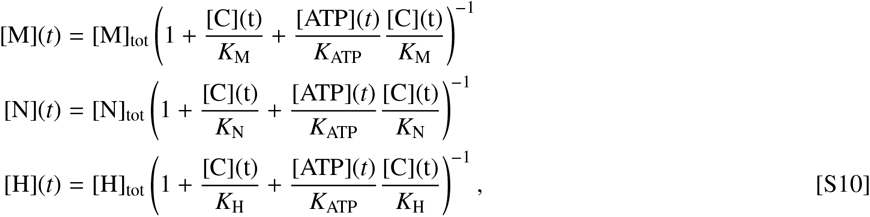

we solve for [C](*t*) in Eq. S9. For the model of CYT-19 activity on pre-RNA, the mass balance relation is given by

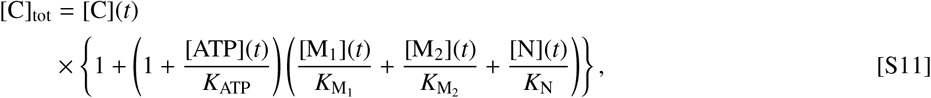

where [M_1_], [M_2_] and [N] are the concentrations of free ribozyme in the M_1_, M_2_, N states, and 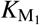, 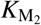 and *K*_N_ denote the respective dissociation constants from CYT-19. Similarly to Eq. S10, the free ribozyme concentration in each state can be expressed as

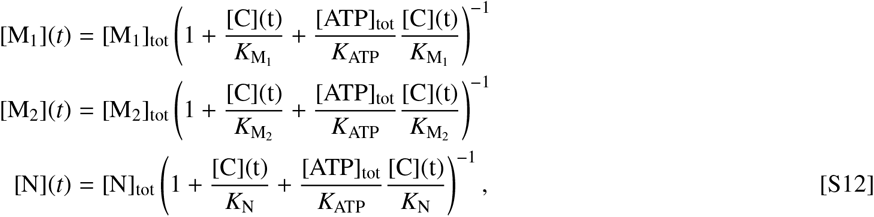

where we keep the ATP concentration, [ATP]_tot_, constant in time.

The IAM-based model in Ref. (23) was developed under the assumption that [C](t) ≈ [C]_tot_. This was feasible in part because the model was fit to the dynamics of RNA folding measured by Bhaskaran *et. al*. (35), in which the experiments were mostly performed with a high CYT-19 concentration relative to the ribozyme concentration; [R]_tot_ = 200 nM and [C]_tot_ ranged from 0 to 3 *μ*M. However, in more recent works by Jarmoskaite *et. al*. (36, 37), the folding of ribozyme was mostly measured in conditions where [C]_tot_ was similar to, or smaller than, [R]_tot_. Thus, to address all the available data on the folding of ribozyme from the three experimental studies (35–37), we take into account the mass balance relation in Eq. S9.

### 3. Mean first passage time

We provide additional context to the thermodynamic cost to refold a misfolded ribozyme to the native state, ⟨*A*_M→N_⟩ (Eq. 10 in the main text). The cost associated with refolding the native ribozyme into the misfolded state, ⟨*A*_N→M_⟩, can be obtained analogously. In order to evaluate the cost of forming the N state from the M state, we set the transition rates out of the N state to 0 (i.e. *k*_NM_ = *k*_cat,NI_ = 0 in Eq. 7). Furthermore, we assume that CYT-19 does not bind productively to the N state (i.e. [C](*t*)/*K*_N_ ≪ 1). Then, the relative concentrations of ribozyme in the I and M states, *P*_I_(*t*) = [I](*t*)/[R]_tot_ and *P*_M_(*t*) = [M](*t*)/[R]_tot_, evolve in time, following the equation,

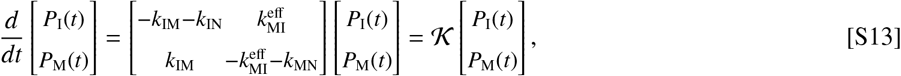

where 𝒦 is a 2 × 2 matrix, and the chaperone-mediated unfolding rate, 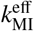, is assumed to be independent of time. The ribozyme eventually transitions to the N state with the reaction rates *k*_IN_ and *k*_MN_, so that the relative concentration of the N state, *P*_N_(*t*) = [N](*t*)/[R]_tot_, increases with rate *dP*_N_(*t*)/*dt* = *k*_IN_*P*_I_(*t*) + *k*_MN_*P*_M_(*t*). With the initial condition *P*_M_(0) = 1, *P*_I_(0) = *P*_N_(0) = 0, the mean first passage time (MFPT) for the ribozyme to transition from the M state to the N state can be written as 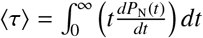, where (*dP*_N_(*t*)/*dt*)|_*t*=*τ*_ is the probability that the ribozyme transitions into the N state at time τ. By using the equality *P*_N_(*t*) = 1 − *P*_M_(*t*) − *P*_I_(*t*) and integration by parts,

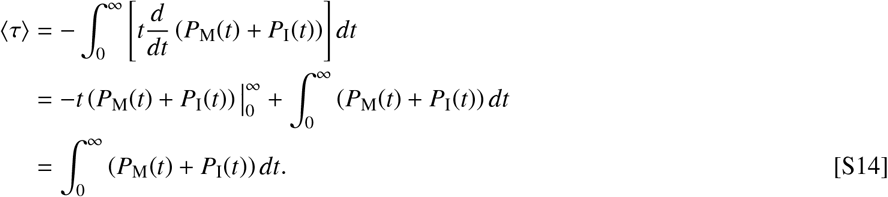

The inverse of MFPT, ⟨τ⟩^−1^, is the rate of N state formation from the M state.

For a given duration 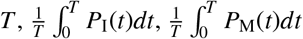, and 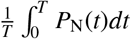 are the time-averaged probabilities for the ribozyme to be found in the I, M and N states, respectively. Thus, the expression 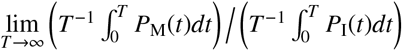 quantifies the ratio between the time-averaged probabilities of finding the ribozyme in the M versus I state. In other words, ⟨τ⟩ can be decomposed into two parts, 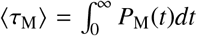 and 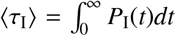, that represent the mean survival times in the M and I states before reaching the N state. With an assumption that only CYT-19 that is bound to the M state hydrolyzes ATP with the time-independent effective rate constant 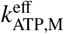, the average ATP cost per refolded ribozyme is given by

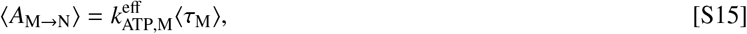

which is equivalent to Eq. 10 in the main text.

To obtain the expression for ⟨τ_M_⟩, we first define the eigenvalues λ_1_ and λ_2_ along with the corresponding eigenvectors **v**_1_ and **v**_2_ of 𝒦 in Eq. S13, so that λ_1_**v**_1_ = 𝒦**v**_1_ and λ_2_**v**_2_ = 𝒦**v**_2_. Next, we define the matrix of eigenvectors **V** = [**v**_1_, **v**_2_]. Then, the solution to Eq. S13 can be written as

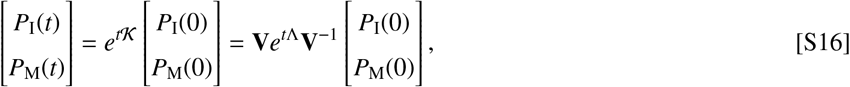

where 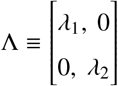. Assuming that λ^1^, λ^2^ < 0, evaluating the integral leads to

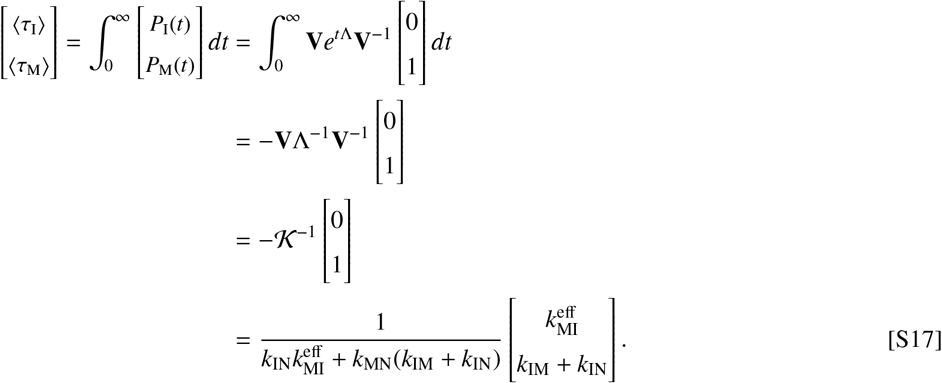

Thus, 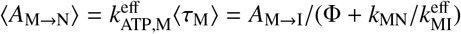, where Φ ≡ *k*_IN_/(*k*_IN_ + *k*_IM_) and 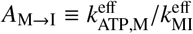.

### 4. Model fit to data

We provide more detail on how the theoretical model (Eq. 7) is fit to the experimental measurements from Refs. (35–37). The quality of fit of the 10-dimensional array of parameters, 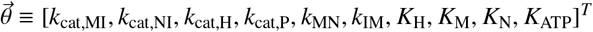 is evaluated by the corresponding χ^2^ value,

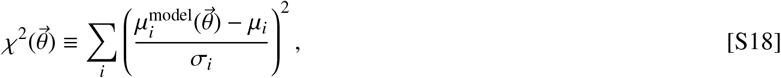

where *i* runs through each measured data point with mean *μ*_*i*_ and standard deviation *σ*_*i*_. 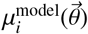 is the simulated data point that matches *μ*_*i*_. The remaining parameters, *k*_IN_ and *k*_NM_, are set by the constraints Φ = 0.1 = *k*_IN_/(*k*_IN_ + *k*_IM_) and *k*_MN_/*k*_NM_ = 10.

To fit the initial rate constants reported in Refs. (36, 37), we consider the initial condition with [M](0) = 200 nM, [I](0) = [N](0) = 0 nM, [C]_tot_ = 1 *μ*M and [ATP]_tot_ = 0.5 mM. First, Eq. 7 is numerically solved while assuming that ATP concentration remains constant in time. The proxy for the time required to reach steady state, *t*^ss^, is defined as the time point at which the inequality 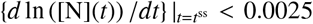 holds, where *t* is in minutes. Then, Eq. 7 is solved a second time from the initial condition until time *t*^ss^. For this second run, ATP concentration, [ATP](*t*), is also modeled as a time dependent variable. The initial rate of N state formation is evaluated at time *t*_init_, defined by [N](*t*_init_) = [N](*t*^ss^)/3. Thus, the initial rate of N state formation is [N](*t*_init_)/*t*_init_, and the initial rate of ATP consumption is ([ATP](0) − [ATP](*t*_init_))/*t*_init_. For the experiments in which all the ribozyme is initially in the N state, *t*_init_ is analogously defined as ([N](*t*_init_) − [N](0)) = ([N](*t*^ss^) − [N](0)) /3.

For most of the measured initial rate constants, both the mean (*μ*) and standard deviation (*σ*) were reported. For the initial rate constants reported without replicate experiments, the standard deviation is set so that the ratio *α* = *μ*/*σ* is equal to twice the average *α* value computed from the rest of the data points belonging to the same family of experiments. For the time course data, the numerical solutions are directly compared with the measurements of the fraction of N state from Ref. (35) (Fig. S1A). No replicate experiments were reported for the time course data, and the standard deviations are set to one-tenth the mean value, so that *σ* = *μ*/10 (see additional file SIdata.pdf for all values used).

### 5. Markov Chain Monte Carlo (MCMC) sampling

We performed MCMC sampling to characterize the distribution of parameters that yield a good fit. To generate the starting point for the MCMC algorithm, each element of the parameter vector is uniformly sampled from the log space with the following bounds: 10^−2^ min^−1^ < *k*_cat,MI_, *k*_cat,NI_ < 10^2^ min^−1^, 10^0^ min^−1^ < *k*_cat,H_, *k*_cat,P_ < 10^3^ min^−1^, 10^0^ min^−1^ < *k*_MN_ < 10^3^ min^−1^, 10^−1^ min^−1^ < *k*_IM_ < 10^2^ min^−1^, 10^−1^ nM < *K*_H_, *K*_M_, *K*_N_ < 10^4^ nM, and 10^2^ *μ*M < *K*_ATP_ < 10^4^ *μ*M. The bounds for *k*_cat,H_, *k*_cat,P_, *k*_MN_, *k*_IM_, and *K*_ATP_ are based on the previous experimental measurements as further detailed in Table. 1. For *k*_cat,MI_ and *k*_cat,NI_, the bounds are set to reflect the range of measured rates for N state formation. For *K*_H_, *K*_M_, and *K*_N_, the bounds reflect the range of CYT-19 concentrations assayed in the experiments. The remaining parameters are set by the constraints Φ = *k*_IN_/(*k*_IN_ + *k*_IM_) = 0.1 and *k*_MN_/*k*_NM_ = 10. By repeated random sampling, we generate the parameter vector 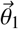 with a sufficiently good fit, 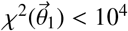, which is used as the starting point of the Metropolis-Hastings algorithm.

For the *i*^th^ iteration of Metropolis-Hastings algorithm, we sample a candidate parameter vector 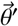^′^ from a multivariate Gaussian centered at 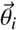, 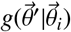. The covariance matrix of 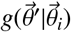, Σ, is a 10 × 10 diagonal matrix, in which each entry is proportional to the range of the corresponding upper and lower bounds used to sample 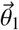. Specifically, for the *j*^th^ parameter 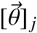, we set [Σ] _*j j*_ = 0.002 (log_10_(*u*_*j*_) − log_10_(*l*_*j*_)), where *u*_*j*_ and *l* _*j*_ are the upper and lower bounds. If the trial parameter yields a better fit (i.e. 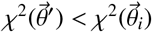), we accept it and set 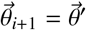. Otherwise, we determine the acceptance by generating a random number *β* ∈ [0, 1], and comparing it with exp (0.5 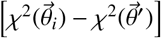^′^). We set 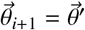 if *β* < exp (0.5 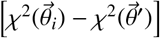) ; otherwise 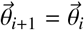 and start the (*i* + 1)^th^ iteration by resampling for the trial vector 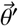.

In total, 400 independent trials of Metropolis-Hastings algorithm were performed, with 3 × 10^5^ iterations each. Out of the 400 trials that started at random initial points, 202 converged to near-optimal χ^2^ values, and generated ∼10^7^ instances of 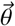 with 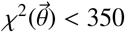. The best fit-parameter array 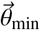, for which 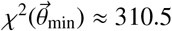, is given in Table 1. The distribution of parameters that yielded relatively small χ^2^ values, as well as the pairwise correlations between the parameters, are shown in Fig. S2.

### 6. Self-splicing yield

The minimal model of self-splicing *in vivo* in the main text describes the time evolutions of the concentrations of pre-RNA in the I, M, N, and SP states, denoted by the column vector 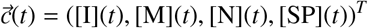. Here, in a slightly more general model than that described in Eq. 12, we assume that the pre-RNA in the intermediate states I, M and N are degraded at the rate *k*_d,i_, and pre-RNA in the spliced state SP are degraded at the rate *k*_d,s_. Then, the pre-RNA molecules evolve with time as

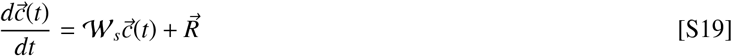

where 𝒲_*s*_ is

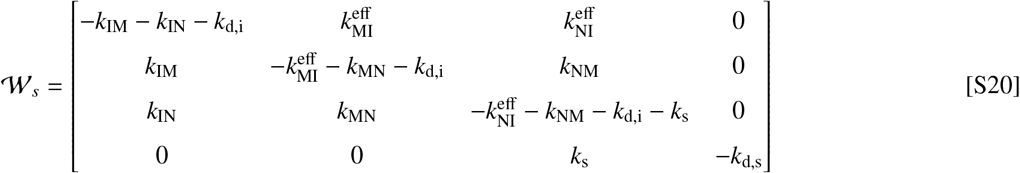

By solving Eq. S19, we obtain the array of steady state concentrations, 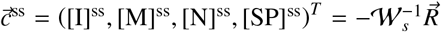. Then, with the total concentration 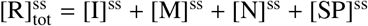, the self-splicing yield is given by

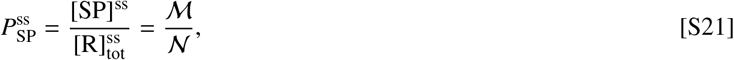

where

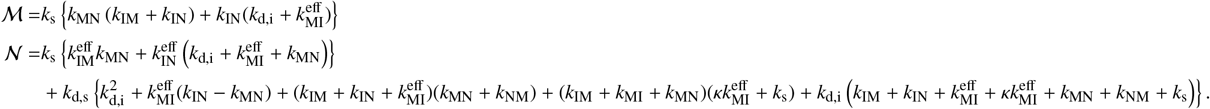

When direct transitions between the M and N states are negligible (*k*_MN_, *k*_NM_ ≪ 1), 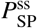 is approximated to

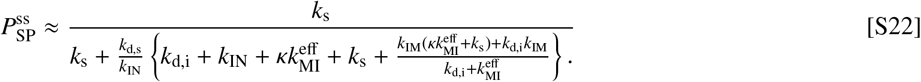

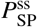 in Eq. S22 is maximized with respect to 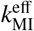

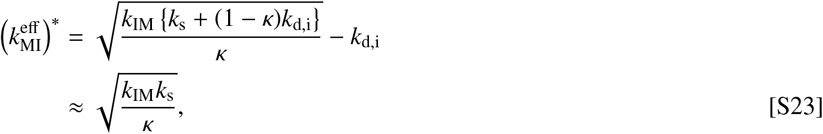

where the approximation holds if *k*_d,i_ ≪ *k*_IM_, *k*_s_. When 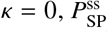 is a hyperbolic function of 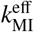, saturating to its maximum value. Qualitatively similar conclusions are drawn for the case with finite values of *k*_MN_ and *k*_NM_ (Fig. 5B).

### 7. Dependence of 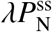 on *κ*

We discuss the *κ*-dependence of 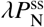 that was used to quantify the steady state yield of native ribozyme over the folding time in Ref.(23). The probability of ribozyme in the I, M, and N states, denoted by the column vector 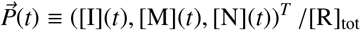, evolve in time following 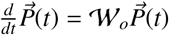, where 𝒲_*o*_ is

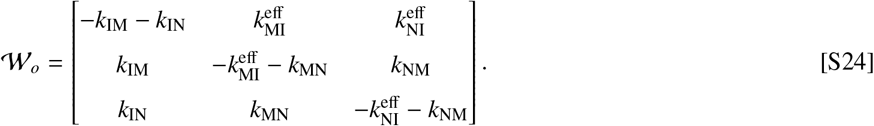

When all the rates are constant in time, 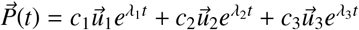, where λ_1_ = 0 > λ_2_ > λ_3_ are the eigenvalues of 𝒲_*o*_, and *u*_1_, *u*_2_, and *u*_3_ are the corresponding eigenvectors. Thus, if |λ_2_| ≪ |λ_3_|, 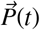 exponentially relaxes to the steady state, with the relaxation rate λ = |λ_2_|. With the initial condition *P*_N_(0) = 0, *P*_N_(*t*) is approximated as

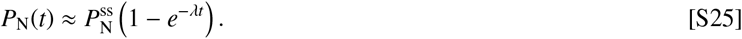

Provided that the transitions between the M and N states are negligible (*k*_MN_, *k*_NM_ ≪ 1), λ and 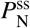 are expressed as follows with 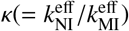 and Φ(= *k* _IN_ / (*k* _IN_+ *k* _IM_)),

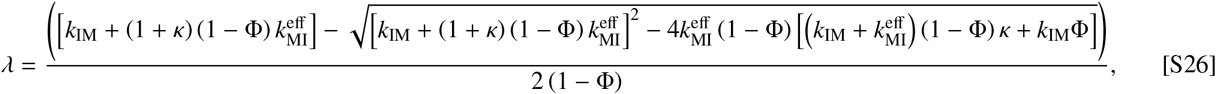

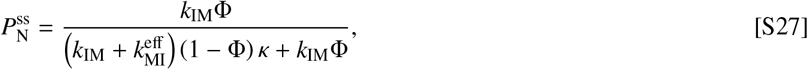

For 0 ≲ Φ ≪ 1 and 0 ≪ Φ ≲ 1, 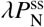 can be expanded as a Taylor series of Φ and 1 − Φ, respectively: For 0 ≲ Φ ≪ 1,

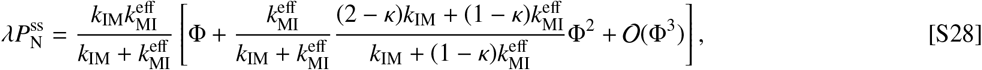

and for 0 ≪ Φ ≲ 1,

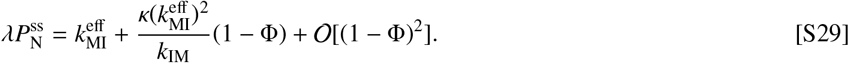

For 0 ≲ Φ ≪ 1, the relative difference between the values of 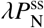 with *κ* = 0 and *κ* = 1 is given as

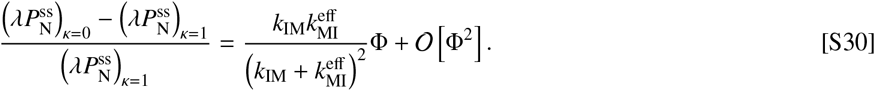

Therefore, the difference between 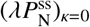 and 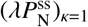 is negligible when 0 ≲ Φ ≪ 1. Qualitatively similar conclusions for the case with finite values of *k*_MN_ and *k*_NM_ are shown in Fig. S3, in which we use the rate constants in Table 1.

**Fig. S1.**
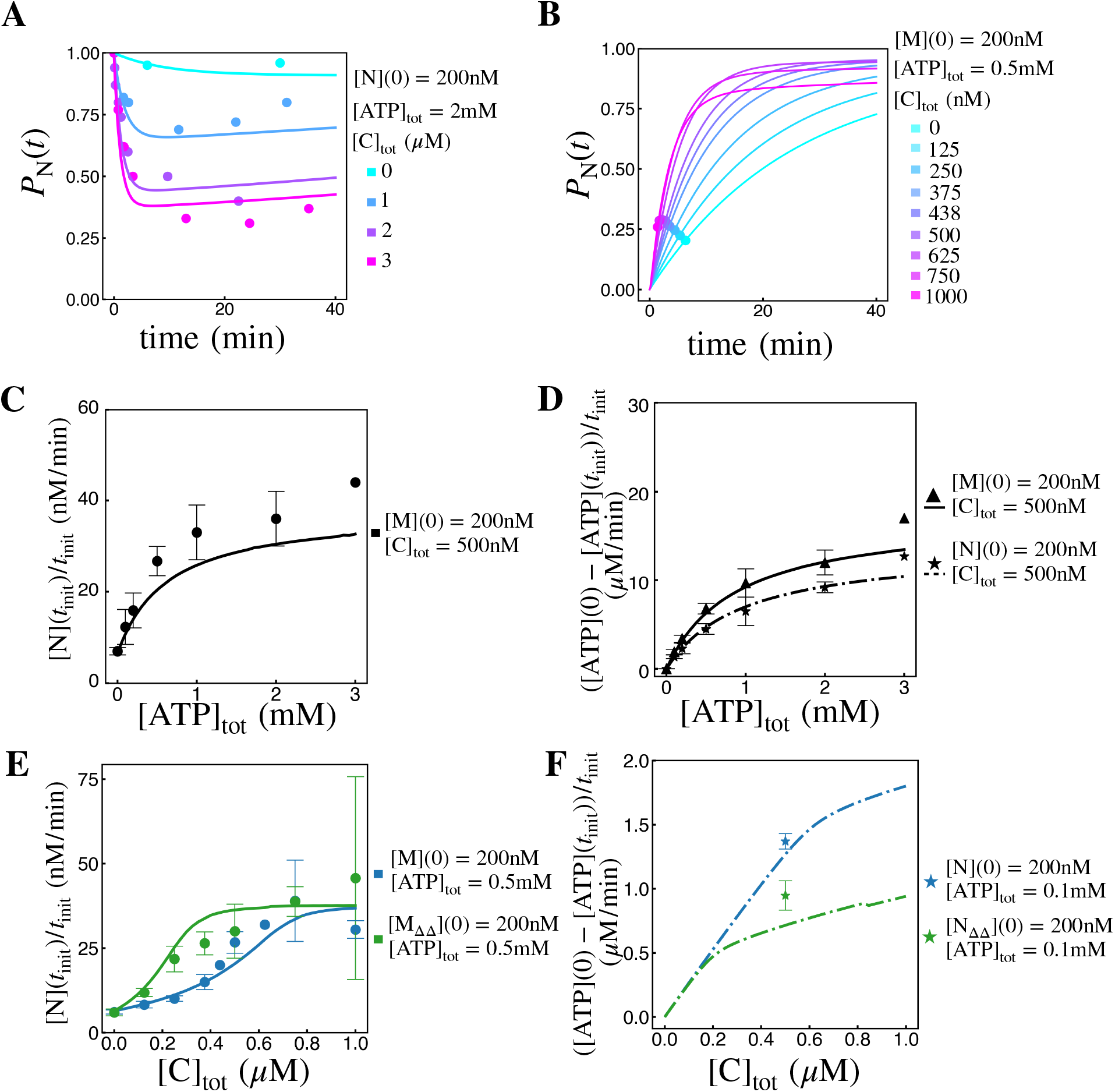
Analysis of CYT-19-facilitated folding of *T*. ribozyme. The curves represent the fitted model, and the dots (except in panel (B)) represent the experimental data. The error bars associated with the dots denote two standard deviations from the mean, and the error bars are omitted for data points generated from single trials. All curves are computed with the best-fit parameters shown in Table 1. **A**. The fraction of N state ribozyme in time (*P*_N_(*t*)) at different concentrations of CYT-19, with the initial condition [N](0) = [R]_tot_ = 200 nM. **B**. The fraction of N state ribozyme (*P*_N_(*t*)) at different concentrations of CYT-19, with the initial condition [M](0) = [R]_tot_ = 200 nM. The filled circles represent the initial time points (*t*_init_) from which the initial rates [N](t_init_)/t_init_ plotted in blue in Fig. 2C are derived. **CD**. The initial refolding rate (C) and ATP hydrolysis rate (D) measured at different concentrations of ATP. **EF**. The initial refolding rate (E) and the initial ATP hydrolysis rate (F) of the mutant ribozyme devoid of P9-2 and P6b (denoted by M_ΔΔ_ and N_ΔΔ_) are shown in green. The Mg^2+^ concentration in each experiment was 2 mM for all but the condition [C]_tot_ = 0 in panel A.

**Fig. S2.**
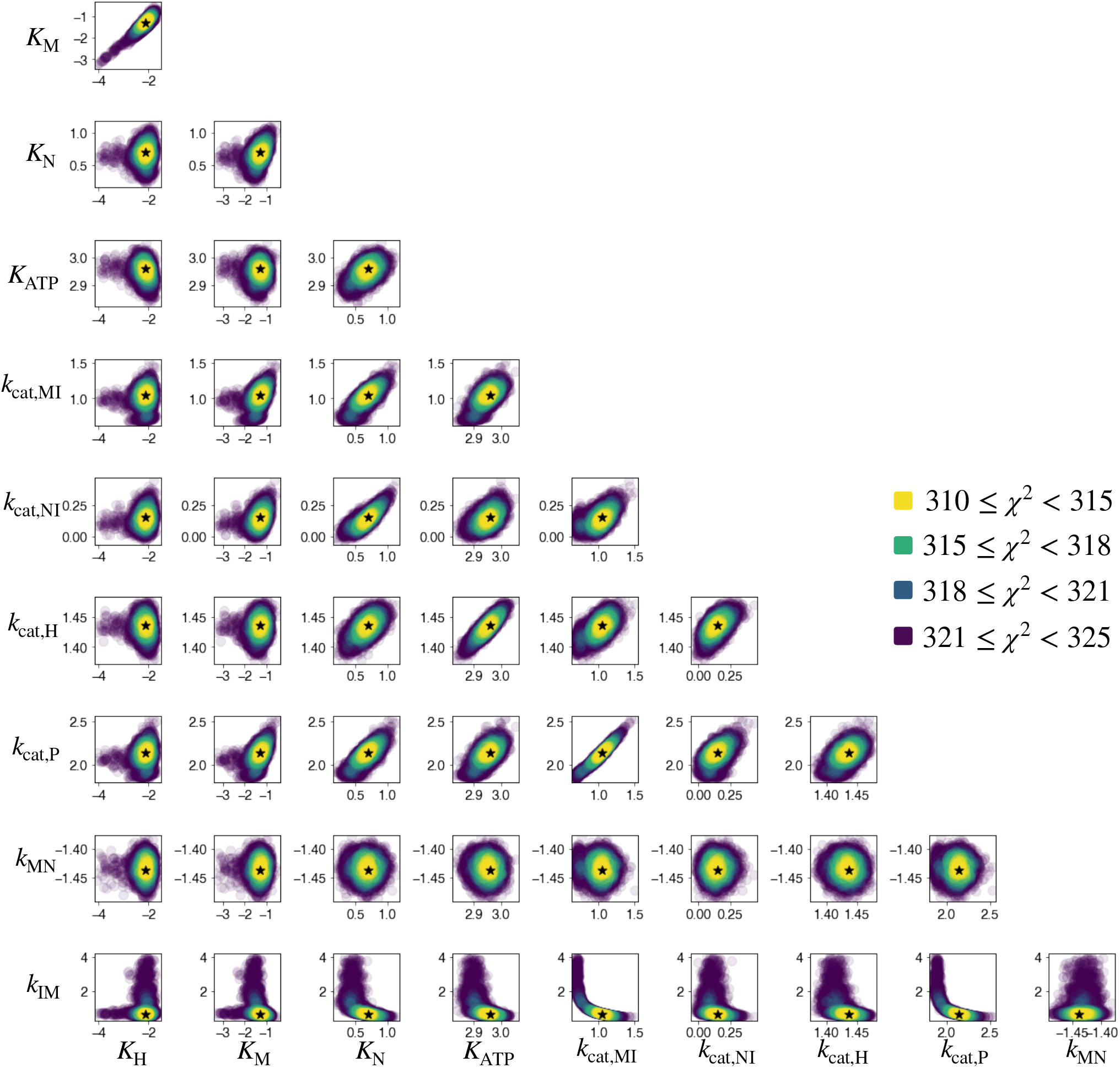
The distribution of parameters that fit experimental data, generated by Monte-Carlo-Markov-Chain (MCMC) method. 400 independent trials of Metropolis-Hastings algorithm were performed to sample the parameters in log space. The pair-wise correlations between all parameters are shown, with the color representing the goodness of fit, 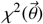 (Eq. S18). The black stars indicate the best-fit parameter vector 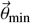. *k*_MN_, *k*_IM_, *k*_cat,MI_, *k*_cat,NI_, *k*_cat,H_, and *k*_cat,P_ are in units of min^−1^, and *K*_H_, *K*_M_, *K*_N_, and *K*_ATP_ are in units of *μ*M. All axes are shown in log_10_ scale.

**Fig. S3.**
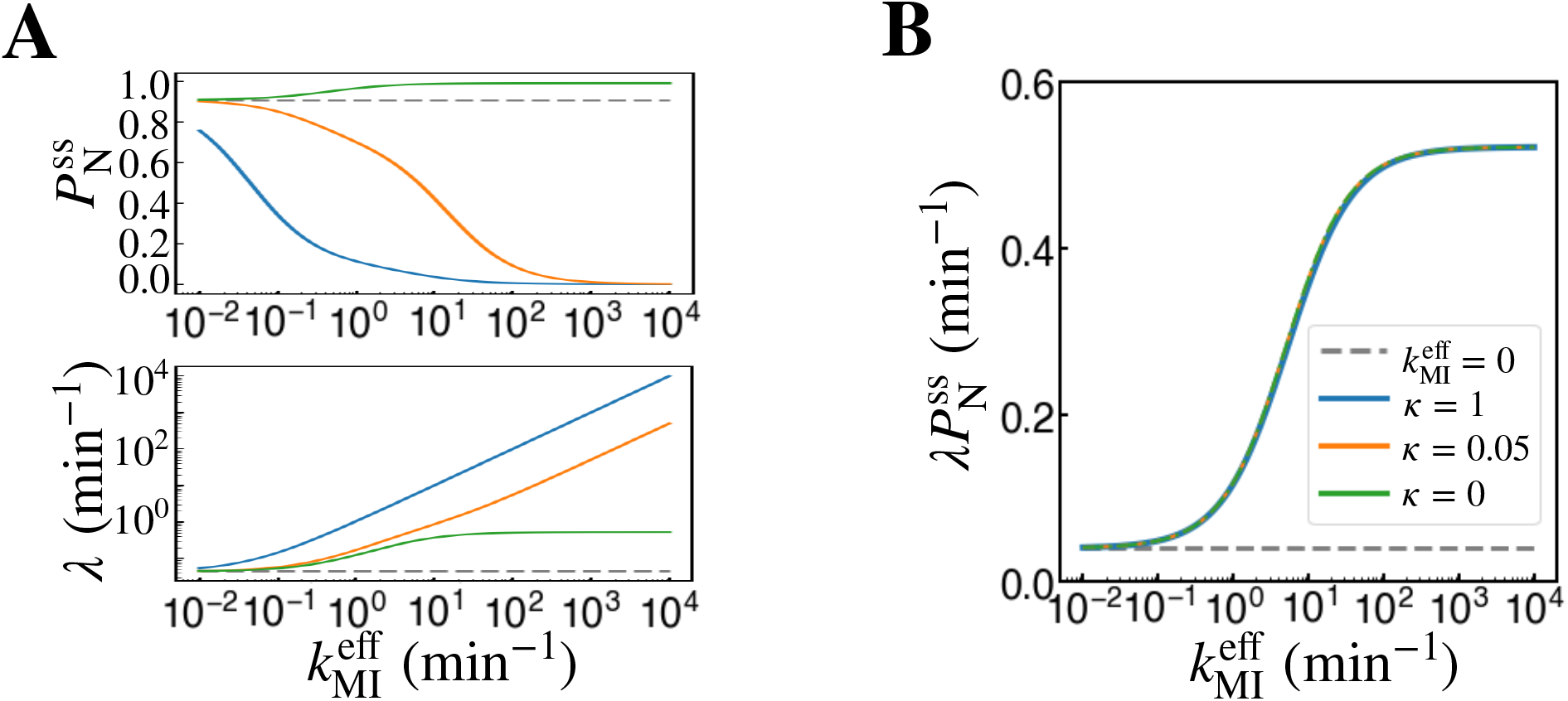
Dependence of 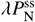 on *κ*. **A** (top) 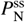 and (bottom) λ, **B** 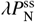 as functions of 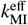, plotted with different *κ* values. The panels **A** and **B** are color-coded identically. The parameters other than 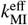 and 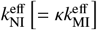 are set to *k*_IM_ = 4.7 min^−1^, *k*_IN_ = 0.53 min^−1^, *k*_MN_ = 0.04 min^−1^, and *k*_NM_ = 0.004 min^−1^.

