## Supplementary material for "Moderate activity of RNA chaperone maximizes the yield of self-spliced pre-RNA *in vivo*": SI data

**Measurements of the initial rate of refolding and ATP hydrolysis**

**Obtained from Jarmoskaite et al., PNAS, 2014 and Jarmoskaite et al., JBC, 2020**

initial condition: 200 nM of M state ribozyme

| CYT-19 (nM) | ATP (mM) | refolding rate (nM/min) | stdev | ATP hydrolysis rate (uM/min) | stdev |
|---|---|---|---|---|---|
| 500 | 0 | 7 | 0.4 | | |
| 500 | 0.1 | 12.3 | 1.9 | 1.9 | 0.15 |
| 500 | 0.2 | 15.9 | 1.9 | 3.4 | 0.2 |
| 500 | 0.5 | 26.7 | 1.6 | 6.8 | 0.3 |
| 500 | 1 | 33 | 3 | 9.7 | 0.8 |
| 500 | 2 | 36 | 3 | 12 | 0.7 |
| 500 | 3 | 44 | 7.3* | 17 | 2.1 |

Initial condition: 200 nM of N state ribozyme

| CYT-19 (nM) | ATP (mM) | | ATP hydrolysis rate (uM/min) | stdev |
|---|---|---|---|---|
| 500 | 0 | | | |
| 500 | 0.1 | | 1.37 | 0.1 |
| 500 | 0.2 | | 2.3 | 0.3 |
| 500 | 0.5 | | 4.5 | 0.3 |
| 500 | 1 | | 6.5 | 0.8 |
| 500 | 2 | | 9.2 | 0.3 |
| 500 | 3 | | 12.7 | 1.7 |

* are obtained from the procedure specified in the SI Appendix section "Model fit to data"

initial condition: 200 nM of M state ribozyme

| CYT-19 (nM) | ATP (mM) | refolding rate (nM/min) | stdev | ATP hydrolysis rate (uM/min) | stdev |
|---|---|---|---|---|---|
| 0 | 0.5 | 6.1 | 0.3 | | |
| 125 | 0.5 | 8.3 | 0.5 | 0.88 | 0.12 |
| 250 | 0.5 | 10.1 | 0.4 | 1.99 | 0.17 |
| 375 | 0.5 | 15 | 1.1 | 4.4 | 0.3 |
| 438 | 0.5 | 20 | 2.3 | 5.2 | 0.2 |
| 500 | 0.5 | 26.7 | 1.6 | 6.8 | 0.3 |
| 625 | 0.5 | 32 | 3.7* | 8.7 | 0.3 |
| 750 | 0.5 | 39 | 6 | 10.7 | 0.7 |
| 1000 | 0.5 | 30.5 | 1.3 | 11.3 | 0.8 |

Initial condition: 200 nM of N state ribozyme

| CYT-19 (nM) | ATP (mM) | | ATP hydrolysis rate (uM/min) | stdev |
|---|---|---|---|---|
| 0 | 0.5 | | | |
| 125 | 0.5 | | 0.76 | 0.1 |
| 250 | 0.5 | | 1.84 | 0.17 |
| 375 | 0.5 | | 3.7 | 0.2 |
| 438 | 0.5 | | 4.5 | 0.3 |
| 500 | 0.5 | | 4.5 | 0.3 |
| 625 | 0.5 | | 5.8 | 0.2 |
| 750 | 0.5 | | 6.9 | 0.4 |
| 1000 | 0.5 | | 7.6 | 1 |

**Measurements of the initial rate of refolding and ATP hydrolysis**
**Obtained from Jarmoskaite et al., PNAS, 2014 and Jarmoskaite et al., JBC, 2020**

initial condition: 200 nM of M state ribozyme

| CYT-19 (nM) | ATP (mM) | refolding rate (nM/min) | stdev | ATP hydrolysis rate (uM/min) | stdev |
|---|---|---|---|---|---|
| 0 | 2 | 7.9 | 0.8 | | |
| 250 | 2 | 15 | 1.6* | 5 | 1.4 |
| 375 | 2 | 22 | 0.5 | 7.3 | 0.9 |
| 500 | 2 | 36 | 3 | 12 | 0.7 |
| 625 | 2 | 42 | 4.5* | 15 | 0.3 |
| 750 | 2 | 45 | 6 | 17.3 | 0.4 |
| 1000 | 2 | 56 | 6.1* | 27.7 | 1.7 |

Initial condition: 200 nM of N state ribozyme

| CYT-19 (nM) | ATP (mM) | | ATP hydrolysis rate (uM/min) | stdev |
|---|---|---|---|---|
| 0 | 2 | | | |
| 250 | 2 | | 5 | 2 |
| 375 | 2 | | 5.4 | 0.6 |
| 500 | 2 | | 9.2 | 0.3 |
| 625 | 2 | | 12 | 0.2 |
| 750 | 2 | | 14 | 2 |
| 1000 | 2 | | 14.3 | 0.4 |

* are obtained from the procedure specified in the SI Appendix section "Model fit to data"

initial condition: 50 nM of M state ribozyme

| CYT-19 (nM) | ATP (mM) | refolding rate (nM/min) | stdev |
|---|---|---|---|
| 0 | 0.5 | 2 | 0.15 |
| 50 | 0.5 | 2.6 | 0.15 |
| 100 | 0.5 | 4 | 0.25 |
| 150 | 0.5 | 9 | 4 |
| 250 | 0.5 | 9 | 0.71 |
| 500 | 0.5 | 9.6 | 0.13 |
| 750 | 0.5 | 9.1 | 5 |

Initial condition: 200 nM M state ribozyme, ATP 100 uM

| ATP hydrolysis rate (uM/min) | stdev |
|---|---|
| 1.37 | 0.06 |

Initial condition: 200 nM M state mutant ribozyme, ATP 100 uM

| ATP hydrolysis rate (uM/min) | stdev |
|---|---|
| 0.95 | 0.11 |

initial condition: 200 nM of M state mutant ribozyme

| CYT-19 (nM) | ATP (mM) | initial refolding rate (nM/min) | stdev |
|---|---|---|---|
| 0 | 0.5 | 6 | 0.5 |
| 125 | 0.5 | 11.9 | 0.6 |
| 250 | 0.5 | 21.8 | 1.9 |
| 375 | 0.5 | 26.5 | 1.7 |
| 500 | 0.5 | 30 | 4 |
| 750 | 0.5 | 38.5 | 2.2 |
| 1000 | 0.5 | 45.6 | 15 |

**Time course experiments with initial concentration of 200 nM of N state ribozyme**

**Obtained from Bhaskaran et al., Nature, 2007**

| concentration of CYT-19=0 | time (min) | Fraction of N state |
|---|---|---|
| | 0 | 1 |
| | 6 | 0.95 |
| | 30 | 0.96 |

| concentration of CYT-19=1 uM | time (min) | Fraction of N state |
|---|---|---|
| | 0 | 1 |
| | 1.7 | 0.82 |
| | 2.6 | 0.8 |
| | 11.7 | 0.69 |
| | 22 | 0.72 |
| | 31.2 | 0.8 |

| concentration of CYT-19=2 uM | time (min) | Fraction of N state |
|---|---|---|
| | 0 | 1 |
| | 0.11 | 0.94 |
| | 0.27 | 0.87 |
| | 0.8 | 0.8 |
| | 1.27 | 0.74 |
| | 2.5 | 0.6 |
| | 9.7 | 0.5 |
| | 22.5 | 0.4 |

| concentration of CYT-19=3 uM | time (min) | Fraction of N state |
|---|---|---|
| | 0 | 1 |
| | 0.75 | 0.77 |
| | 1.8 | 0.62 |
| | 3.4 | 0.5 |
| | 13 | 0.33 |
| | 24.5 | 0.31 |
| | 35.2 | 0.37 |
